# Protein embeddings and deep learning predict binding residues for various ligand classes

**DOI:** 10.1101/2021.09.03.458869

**Authors:** Maria Littmann, Michael Heinzinger, Christian Dallago, Konstantin Weissenow, Burkhard Rost

## Abstract

One important aspect of protein function is the binding of proteins to ligands, including small molecules, metal ions, and macromolecules such as DNA or RNA. Despite decades of experimental progress many binding sites remain obscure. Here, we proposed *bindEmbed21*, a method predicting whether a protein residue binds to metal ions, nucleic acids, or small molecules. The Artificial Intelligence (AI)-based method exclusively uses embeddings from the Transformer-based protein Language Model (pLM) ProtT5 as input. Using only single sequences without creating multiple sequence alignments (MSAs), *bindEmbed21DL* outperformed MSA-based predictions. Combination with homology-based inference increased performance to F1=48±3% (95% CI) and MCC=0.46±0.04 when merging all three ligand classes into one. All results were confirmed by three independent data sets. Focusing on very reliably predicted residues could complement experimental evidence: For the 25% most strongly predicted binding residues, at least 73% were correctly predicted even when ignoring the problem of missing experimental annotations. The new method *bindEmbed21* is fast, simple, and broadly applicable - neither using structure nor MSAs. Thereby, it found binding residues in over 42% of all human proteins not otherwise implied in binding and predicted about 6% of all residues as binding to metal ions, nucleic acids, or small molecules.

## Introduction

### Experimental data for protein binding remains limited

Knowing protein function is crucial to understand the molecular mechanisms of life^1^. For most proteins, function depends on binding to other molecules called *ligands*^2^; these include metal ions, inorganic molecules, small organic molecules, or large biomolecules such as DNA, RNA, and other proteins. Although the variation in binding sites resembles the diversity of the ligands, binding sites are highly specific and often determined by a few key residues^2^. Binding residues are experimentally determined most reliably through high-resolution structures of the protein in complex with the ligand marking residues close to this ligand as binding residues (e.g., ≤ 5Å)^3^.

### Prediction methods usually rely on MSAs

Despite immense progress in quantitative high-throughput proteomics, experimentally verified binding residues remain unknown for most proteins^4^. In fact, reliable data remain so sparse to even challenge Machine Learning (<u>ML</u>) models with fewer parameters than tools from Artificial Intelligence (<u>AI</u>)^5^. Thus, reliable prediction methods importantly bridge, e.g., studying the effect of sequence variation in human populations^6,7^. Homology-based inference (<u>HBI</u>) transfers e.g., binding residues from sequence-similar proteins with known annotations to uncharacterized proteins^5,8^. Although accurate, HBI is only applicable to the few proteins for which a sequence-similar protein with binding annotations exists. If unavailable, *de novo* prediction methods based on ML try to fill the gap. HBI (or template-based methods) usually outperforms sequence-based (or *de novo*) methods^9,10^, but they rely on the existence of structurally similar proteins with experimentally verified binding annotations (structural template)^10–14^. Notably, *AlphaFold 2* that solved the protein structure prediction problem^15^ might be the first AI-based prediction method consistently outperforming template-based solutions. AlphaFold 2 heavily relies on information from multiple sequence alignments (<u>MSAs</u>). Recent structure predictions without MSAs remain less accurate^16^. It remains unclear to which extent structure predictions could improve binding prediction beyond stepping up from binding residues to binding sites.

Some template-based methods also require substantial computing resources, e.g., COACH^10^ is an ensemble classifier combining five individual approaches and has been considered the state-of-the-art (_SOTA_) for binding residue prediction for many years^17,18^. One protein prediction took about 10 hours on their webserver, while local installations require 60GB free disk space. Although neither aspect renders the method unusable, both limit ease of access to predictions and comparisons. On the other hand, sequence-based methods usually depend on sufficiently diverse and reliable experimental data and expert-crafted input features including evolutionary information to represent protein sequences^5,15,17,19,20^. Our previous method bindPredictML17^5^ predicted binding residues for enzymes and DNA-binding proteins relying mainly on information from sequence variation^21,22^ and co-evolving residues^23^, both requiring the time-consuming computation of MSAs. Another method, ProNA2020^19^, uses MSAs and various features from PredictProtein^24^ to predict protein-protein, protein-DNA, and protein-RNA binding. In addition to the complexity of their input features, many methods specialize on specific ligands or sets thereof^5,14,18–20,25–27^. For instance, PredZinc^20^ only predicts zinc ions and IonCom^18^ provides predictions for 13 metals and for four radical ion ligands. Most existing sufficiently reliable sequence-based methods cannot be applied to generic proteome-wide binding predictions due to restrictions in computational resources or to limited sets of ligands.

Here, we propose a new method dubbed *bindEmbed21* consisting of two components (bindEmbed21DL and bindEmbed21HBI) that predict binding residues for three ligand classes. We input protein representations (fixed-length per-protein embeddings) from pre-trained protein Language Models (<u>pLMs</u>), in particular from ProtT5^28^. Using only those embeddings, bindEmbed21DL predicts residues binding to metal ions, nucleic acids (DNA and RNA), and/or regular small molecules. Combining the *de novo* prediction method with HBI (bindEmbed21HBI) further improved performance. Since embeddings can be easily extracted for any protein sequence, bindEmbed21 enables fast and easy predictions for all available protein sequences.

## Results & Discussion

### Embedding-based predictions from bindEmbed21DL achieved F1=43%

Inputting raw ProtT5^28^ embeddings into a shallow two-layer CNN, our new method, *bindEmbed21DL*, predicted for each residue in a protein, whether or not it binds to a metal ion, a nucleic acid (DNA or RNA), or a small molecule. The prediction performance differed substantially between the three classes (Fig. 1, Table S1 in Supporting Online Material (SOM)): binding residues were predicted best (e.g., highest F1 or MCC, Eqns. 3&4) for small molecules and worst for nucleic acids (Table 1 DevSet1014; Fig. 1A-C). Those differences might point to differences in the abundance of experimental data for each ligand class: Small molecules were the most prominent ligand class, while nucleic binding was the lowest (Table S11). This might suggest that any class could be predicted better given more data. In fact, using a smaller training set (515 proteins) with equal numbers of proteins with small molecule as with nucleic acid binding (108 proteins) dropped performance immensely for the small molecule class (Table S3) suggesting that better prediction of small molecule binding resulted largely from access to more experimental data. Alternatively, performance differences could be due to properties of small molecule binding being more clearly encoded in the embeddings than those of nucleic acid binding. This might render these easier to predict. However, we could neither support this speculation by explicit evidence, nor refute it and proof that only the increase in data caused better predictions. Performance appeared highest when dropping the distinction between ligand classes, i.e., simplifying the task to the prediction of binding vs. non-binding (Table 1; Fig. 1D), indicating that many residues were correctly identified as binding residues despite confusing the ligand classes (Table S5).

**Fig. 1:**
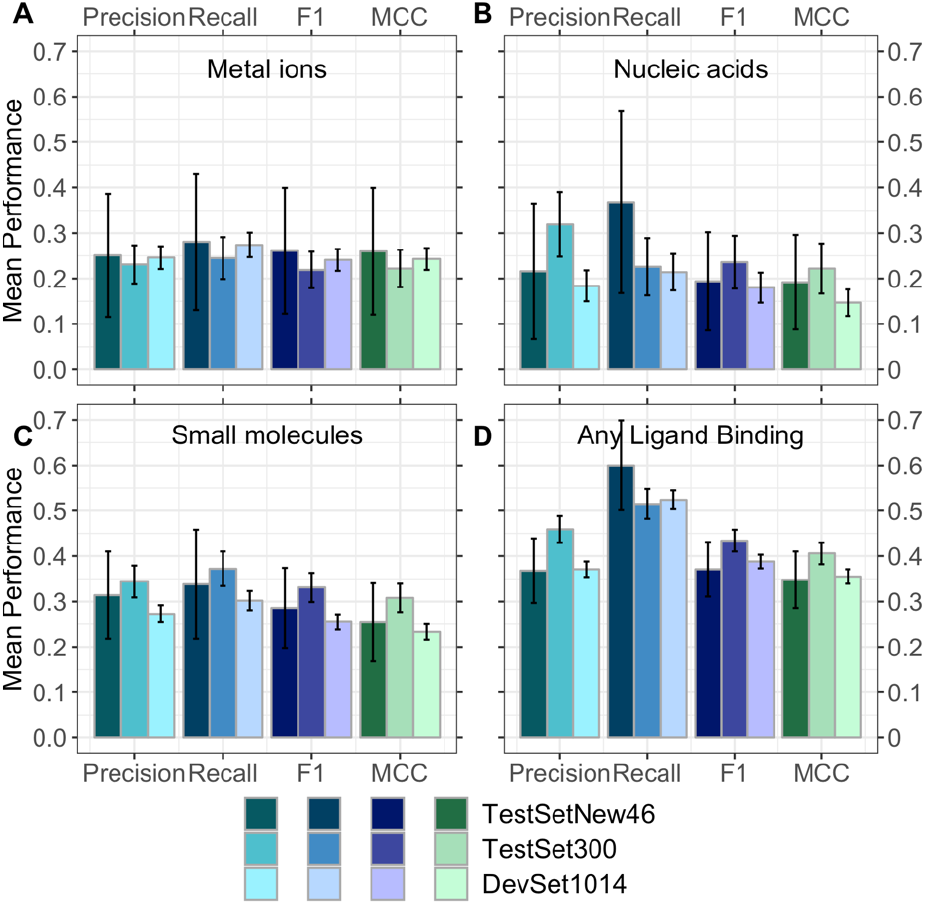
Performance of new method *bindEmbed21DL*. Performance captured by four per-residue measures: precision (Eqn. 2), recall (Eqn. 1), F1 score (Eqn. 3), and MCC (Eqn. 4). <u>Data sets:</u> *DevSet1014* (validation splits of cross-validation development, most light colors), *TestSet300* (fixed test set used during development, darker colors), and *TestSetNew46* (additional test set compiled after development, most dark colors). <u>Predictions of residues binding to</u> **A.** metal ions, **B.** nucleic acids (DNA or RNA), **C.** small molecules, and **D.** any ligand class grouping all three classes into one (considering each residue predicted/observed to bind to one of the three ligand classes as binding, all others as non-binding). On the validation set *DevSet1014, bindEmbed21DL* predicted any binding residue with F1=39±2%. Surprisingly, the number was slightly higher for the test set *TestSet300* (F1=43±2%) while being similar on the additional test set *TestSetNew46* (F1=37±6%). Error bars indicate 95% CIs.

**Table 1:** F1 score (harmonic mean of precision and recall). *.

| Method | Dataset | F1-metal | F1-XNA | F1-small | F1-all |
| --- | --- | --- | --- | --- | --- |
| *bindEmbed21DL* | *DevSet1014* | 24± 2% | 18± 3% | 26± 2% | 39± 2% |
| *bindEmbed21DL* | *TestSet300* | 22± 4% | 24± 6% | 33± 3% | 43± 2% |
| *bindEmbed21DL* | *TestSetNew46* | 26±14% | 19±11% | 29± 9% | 37± 6% |
| *Random* | *TestSet300* | 1± 1% | 6± 2% | 6± 1% | 9± 1% |
| *bindEmbed21DL* | *TestSet225* | n/a | n/a | n/a | 47± 2% |
| *bindPredictML17* | *TestSet225* | n/a | n/a | n/a | 34± 2% |
| *bindEmbed21DL* | *TestSet300<sub>XNA66</sub>* | n/a | 31± 5% | n/a | n/a |
| *ProNA2020* | *TestSet300<sub>XNA66</sub>* | n/a | 33± 7% | n/a | n/a |
| *bindEmbed21DL* | *TestSet300<sub>Zinc51</sub>* | 58± 8% | n/a | n/a | n/a |
| *PredZinc* | *TestSet300<sub>Zinc51</sub>* | 58±10% | n/a | n/a | n/a |
| *ZincBindPredict* | *TestSet300<sub>Zinc51</sub>* | 17± 9% | n/a | n/a | n/a |

\* **Measure:** F1 (Eqn. 3); ± : 95% confidence intervals (1.96 standard errors); **Methods:** *bindEmbed21DL*: method introduced here, *bindPredictML17*<sup>5</sup>: MSA-based method predicting binding, *ProNA2020*<sup>19</sup>: method specialized on predicting binding to DNA, RNA, and other proteins; *PredZinc*<sup>20</sup> and *ZincBindPredict*<sup>29</sup>: methods specialized on predicting zinc-binding; *Random*: random prediction by randomly shuffling the original output probabilities
of `bindEmbed21DL`; **Data:** *DevSet1014*: development set (validation) set with 1,014 proteins, *TestSet300*: Test set created during method development with 300 proteins, *TestSet225*: subset of test set shared with `bindPredictML17`, *TestSetNew46*: 46 sequence-unique proteins added since development of this work began – all sequence-unique with respect to each other and all other proteins used, *TestSet300<sub>XNA66</sub>*: subset with DNA or RNA (dubbed XNA) binding proteins from our test set. *TestSet300<sub>Zinc51</sub>*: subset with zinc-binding proteins from our test set.

For all ligand classes, precision (Eqn. 2) remained below recall (Eqn. 1; Fig. 1) and the fraction of proteins for which not a single residue was predicted as binding (CovNoBind(l), Eqn. 9), was low, especially for metal ions and small molecules (Table S4). Therefore, performance for the individual ligand classes appeared limited by over-prediction (binding predictions not experimentally confirmed, yet) and cross-predictions (predicted to bind ligand C1, annotated for C2). As the binary prediction (binding/not) outperformed by far the 3-class prediction (Table 1), cross-predictions (confusions between ligand classes, Table S5) constituted one major limitation. The most common cause for prediction mistakes appeared to be over-prediction (Table S4), but at least some of the alleged over-predictions might indicate missing observations (analysis below). Remarkably, bindEmbed21DL performed similar to its binarized version solely trained on the distinction of binding vs. non-binding (Table S6).

In a typical cross-validation split (training, validation, test), performance values are higher for the validation than for the test set, because hyper-parameters are optimized on the former. We observed the inverse except for binding to metal ions (Table 1, Fig. 1) although most differences were within the confidence intervals (CI; Fig. 1, Table S1). The test set had more proteins binding to nucleic acids and small molecules than the development set, due to constraints imposed on the test set to facilitate comparisons with other methods. Those were the classes for which bindEmbed21DL reached higher values on the test than on the validation set (Fig. 1B&C). Thus, the higher numbers for the test set for nucleic acid and small molecule binding could indicate that binding residues are better defined and therefore easier to predict for enzymes than for other proteins in the development set.

To investigate, we created an independent test set from recent annotations (TestSetNew46, Methods: 46 unique from a total of 1,592 new proteins). For these, *bindEmbed21DL* reached values that, within the 95% CI, agreed with both the original test and validation sets, possibly due to large CIs for the tiny new data set. When merging all ligand classes, the new test set was large enough to establish with statistical significance (95% CI) that our performance estimates reflected what is to be expected for the next 1,592 proteins submitted for prediction (Methods).

### Embeddings clearly outperformed MSA-based predictions

One recent binding method, *bindPredictML17*^5^ predicts binding residues based on MSAs. A subset of the test set (225 of the 300 proteins in TestSet300) enabled an unbiased comparison: *bindEmbed21DL* significantly (beyond 95% CI) outperformed the old MSA-based method *bindPredictML17*, e.g., raising the harmonic mean over precision and recall by 13 percentage points (Fig. 2, Table 1, bindEmbed21DL vs. bindPredictML17 last column for TestSet225). However, *bindEmbed21DL* predicted binding for only 222 of the 225 test proteins (CovOneBind=99%, Eqn. 8), while its predecessor predicted for all 225. This could be attributed to bindEmbed21DL focusing more on precision than bindPredictML17: The gain in precision was larger than the gain in recall (Fig. 2). However, higher precision reduced recall, thereby missing binding in three of 225 proteins.

**Fig. 2:**
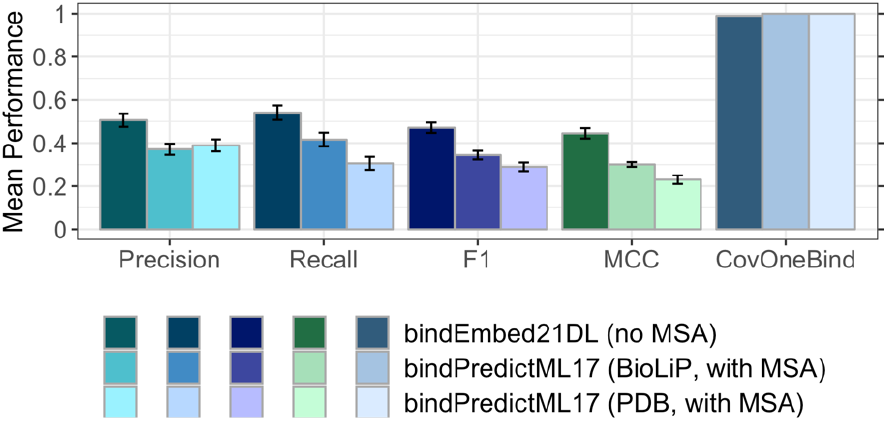
Embeddings outperformed MSA-based predictions. Comparison of performance between bindPredictML175 using multiple sequence alignments (MSAs) and the method introduced here, bindEmbed21DL, using only embeddings from ProtT528. We also compare using binding annotations from BioLiP9 or the PDB30. bindEmbed21DL (embeddings-only) clearly outperformed bindPredictML17 (MSA+BioLiP) by 13 percentage points (F1 =47±2% vs. F1 =34±2%). We used annotations from BioLiP9 to assess the performance for both methods. Although bindPredictML17 had been trained on annotations from PDB30 for enzymes and PDIdb31 for DNA-binding proteins, it reached higher performance (lighter shaded colors vs. lightest shaded colors) for BioLiP annotations. Error bars indicate 95% CIs.

*bindEmbed21DL* and *bindPredictML17* differed in two major aspects: (1) the annotations used for training, and (2) the usage of embeddings vs. MSA-derived input features. Both factors contributed to the improvement of bindEmbed21DL over bindPredictML17. For instance, the F1 score improved by 18 percentage points; 13 of the 18 originated from using embeddings rather than MSA-based input (Fig. S2, SOM 1.3), while five of 18 reflected the new annotations (Fig. 2, SOM 1.2). Thus, embeddings can significantly outperform methods explicitly using evolutionary information through MSAs.

### bindEmbed21DL competitive to specialist methods

*bindEmbed21DL* predicted three ligand classes, while many state-of-the-art (SOTA) methods specialize on one ligand class or subsets thereof. For instance, *ProNA2020*^19^ focuses on predicting protein-, DNA-, or RNA-binding, both on the per-protein (does protein bind DNA or not?) and the per-residue (which residue binds DNA?) level. The MSA-based method ProNA2020 shines through unifying a hierarchy of prediction tasks and outperformed all other sequence-based methods in predicting binding to DNA or RNA (dubbed XNA)^19^. We compared the specialist *ProNA2020* with the generalist *bindEmbed21DL* using 66 nucleic acid binding proteins in *TestSet300* (dubbed TestSet300_xNA66_ in Table 1). For those 66, *ProNA2020* performed slightly worse in XNA-binding prediction than the embedding-based MSA-free *bindEmbed21DL* (Fig. 3A). However, when analyzing how many proteins had at least one residue predicted as XNA-binding (CovOneBind, Eqn. 8), the situation reversed (Fig. 3A). When considering all residues predicted by *bindEmbed21DL* as binding (bind=nucleic acids + metal ions + small molecules), F1 rose almost ten percentage points to 43±5% and CovOneBind to 97% (Fig. 3A). This again pointed to the problem of cross-predictions (Table S5).

**Fig. 3:**
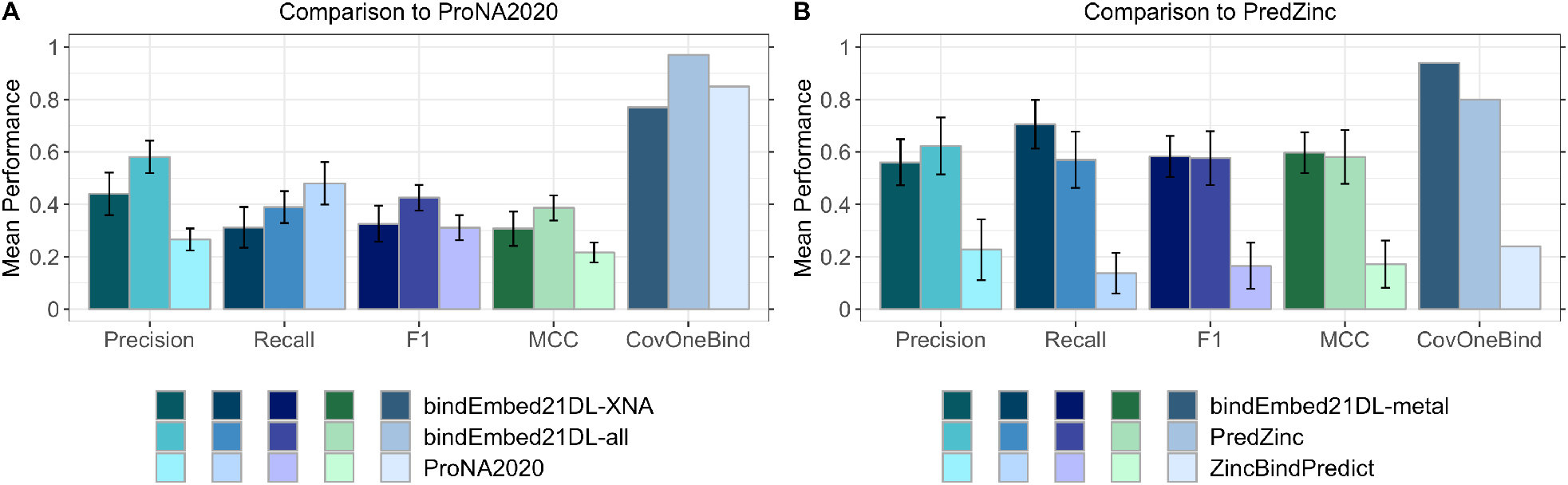
bindEmbed21DL competitive with specialists. Panel A: XNA binding. <u>Data</u>: 66 DNA- or RNA-binding (dubbed XNA) proteins from the test set *TestSet300*. <u>ProNA2020</u>^19^ (lightest shaded bars) uses MSAs to predict DNA-, RNA-, and protein-binding, while the method introduced here uses embeddings only (no MSA); <u>bindEmbed21DL-XNA</u> (darkest shaded bars) marked predictions of either DNA or RNA (XNA); <u>bindEmbed21DL-all</u> (lighter shaded bars) marked using all binding predictions and assessing only XNA-binding. While the difference in F1 scores between the three methods was within the error bars (95% CIs), bindEmbed21DL (-XNA and -all) achieved a statistically significant higher performance than ProNA2020 while ProNA2020 achieved a higher recall. Also, the fraction of proteins with at least one XNA prediction (CovOneBind, Eqn. 8) was higher for ProNA2020 than for bindEmbed21DL-XNA. However, when considering any residue predicted as binding (*bindEmbed21DL-all:* nucleic acid, or metal ion, or small molecule), our new method apparently reached the highest values due to confusions between XNA and other ligands (Table S5). **Panel B: Zinc-binding.** <u>Data</u>: 51 zinc-binding proteins from *TestSet300*. <u>ZincBindPredict</u>^29^ (lightest shaded bars) and _PredZinc_^20^ (darker shaded bars) predict zinc-binding; <u>bindEmbed21DL-metal</u> (darkest shaded bars) marked predictions for metal ions. bindEmbed21DL-metal achieved a similar performance as PredZinc, while providing predictions for more proteins (CovOneBind(bindEmbed21DL-metal)=94% vs. CovOneBind(PredZinc)=80%). ZincBindPredict was not competitive due to only providing predictions for 12 proteins (CovOneBind(ZincBindPredict)=24%).

<u>*PredZinc*</u>^20^ and <u>*ZincBindPredict*</u>^29^ specialize on predicting residues binding to zinc ions. 51 proteins in *TestSet300* were annotated as zinc-binding (dubbed TestSet300_zinc51_ in Table 1) and were used to compare *PredZinc* and *ZincBindPredict* to the generalist *bindEmbed21DL*. The newer method, *ZincBindPredict*, only predicted for 12 proteins (CovOneBind=24%, Fig. 3B). Therefore, we also compared to the older method *PredZinc*. Despite having only been trained on metal-binding in general and not zinc-binding specifically, *bindEmbed21DL* matched the F1 score of *PredZinc* (Fig. 3B) with a lower precision but higher recall (Fig. 3B); it also reached a higher CovOneBind (Eqn. 8) predicting for 94% instead of for 80% as *PredZinc*. Also, *bindEmbed21DL* clearly outperformed *ZincBindPredict* (Fig. 3B) through higher CovOneBind, i.e., *ZincBindPredict* is very accurate when applicable.

We evaluated specialized methods only on proteins binding to those ligand classes. In a more realistic application not knowing the ligand, specialized methods likely perform worse. Also, we may have overestimated the performance of other methods because we could not exclude their development sets. Nevertheless, *bindEmbed21DL* remained competitive on the turf of the specialists and generic enough to be applicable to three different ligand classes.

### More reliable predictions better

For the binary prediction of binding vs. non-binding residues, *bindEmbed21DL* reached 37±2% precision at 52±2%recall (Fig. 1D) while making predictions for 1,000 of 1,014 proteins in the validation splits (*DevSet1014*; CovOneBind=99%). These values resulted from the default threshold (p≥0.5) optimized by the ML method. If only the 1,000 proteins with a prediction were considered, both precision and recall rose by one percentage point (Fig. 4). We analyzed the trade-off between precision, recall, and CovOneBind in dependence of the output probability: Precision decreased for lower cutoffs but recall and CovOneBind increased allowing more binding predictions for more proteins (Fig. 4, Table S7). For instance, at a cutoff of 0.28, at least one binding residue was predicted for every protein (CovOneBind=100%) at the expense of precision dropping by nine percentage points (Fig. 4, Table S7). On the other hand, precision could be increased by applying higher cutoffs to predict binding. For instance, for a cutoff of 0.95, precision almost doubled (Fig. 4, Table S7). Although recall and CovOneBind decreased for higher cutoffs, *bindEmbed21DL* still predicted binding for over half of the proteins and for one fourth of all binding residues at 0.95 (Fig. 4, Table S7). Residues falsely predicted as binding at such a high cutoff could point to yet unknown candidates for binding residues. In fact, comparing the internal representations from the first CNN layer of falsely predicted binding residues with those of correct predictions provided some evidence that highly reliable, not yet observed predictions clustered with those of experimental annotations (Fig. S5). This confirmed the hypothesis that highly reliable binding predictions might help to identify missing binding annotations.

**Fig. 4:**
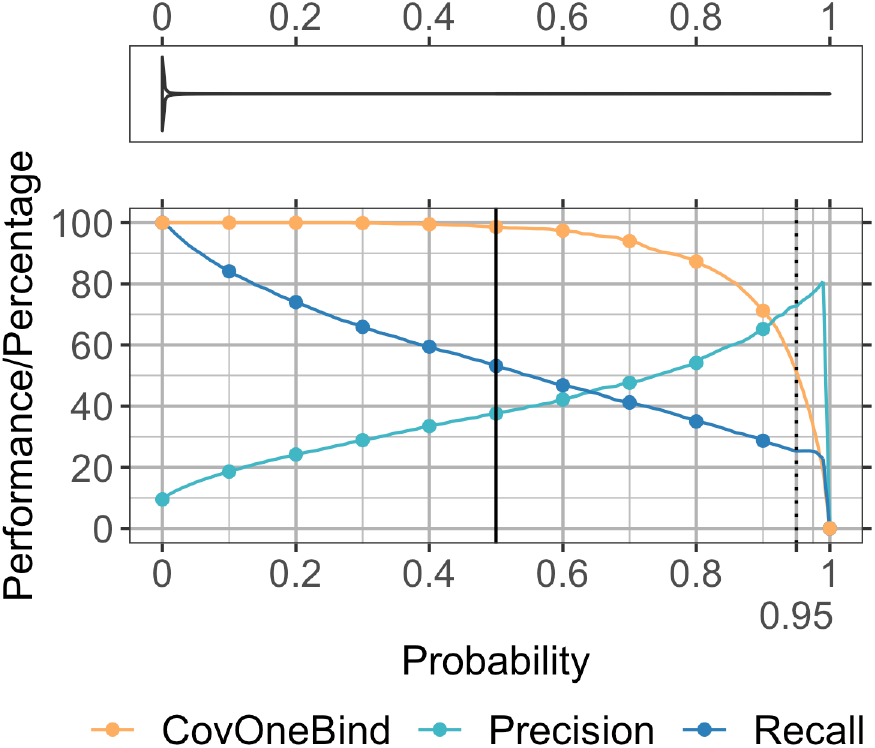
Residues predicted stronger more often correctly predicted. <u>Data set</u>: DevSet1014. Precision and recall are only shown for the proteins for which at least one residue was predicted as binding where the number of such proteins is indicated by CovOneBind. The x-axis gives the output probability of bindEmbed21DL for a prediction corresponding to the prediction strength. The y-axis gives the average performance or percentage of proteins with a prediction at the respective probability cutoff. All curves give the cumulative values, e.g., the precision of all residues predicted with probability ≥ 0.95 (marked as dashed line) was 73% corresponding to a recall of 25%; and at that value, at least one binding residue was predicted in 51% of the proteins. While higher probabilities correspond to more reliable binding predictions, lower probabilities correspond to highly reliable non-binding predictions (Table S7; SOM 1.5 for more details). The violin plot in the top panel reflects the actual distribution of probabilities: 50% of the residues were predicted with probability≤10^−3^ and 75% with probability≤6*10^−3^. While we expected binding to be the more evolutionary conserved feature, non-binding residues were apparently easier to predict reliably.

Missing experimental annotations limit the top precision reachable (if we equate “not observed as binding” with “non-binding”). For probability>0.95, precision reached 80% (Fig. 4). To some extent, this estimated the effect from missing annotations: At least for the 25% most reliably predicted binding residues, at most one fifth (100-80) could be attributed to missing annotations. This might not imply that, at p>0.5, precision would maximally rise by 20 percentage points because the most reliably predicted binding residues might be more likely to coincide with easy to obtain experimental data. Clearly, the opposite holds: Regions with low information, such as intrinsically disordered regions, are more difficult to predict and to experimentally resolve^32^.

Alternatively, predictions could be refined taking the number of predicted residues into consideration: A low number of binding predictions in a protein indicated that those predictions were incorrect (Fig. S6). Removing such predictions led to an increase in CovNoBind(l) (Eqn. 9) while decreasing CovOneBind (Eqn. 8; Fig. S6).

Since the probability score correlated with prediction reliability, we defined a single-digit integer reliability index (RI; Eqn. 10) ranging from 0 (unreliable; probability=0.5) to 9 (very reliable). This RI empowers users, depending on their interest, either to focus on the most precise/reliable predictions for binding (or non-binding), or to focus on the perspective most likely to identify any binding residue that might exist.

### Reliable predictions could help refining experimental annotations

Using a cutoff of 0.95 to classify a residue as “binding”, *bindEmbed21DL* reached 73% precision with at least one residue predicted as binding for 519 proteins (CovOneBind=51%; Fig. 4, Table S7). For 84 of the 519 proteins (16%), none of the residues predicted that reliably (probability≥0.95) had been experimentally annotated as binding. We analyzed two of those 84 in more detail.

The DNA-binding protein HMf-2 (UniProt ID: P19267) is annotated to bind metal with residues 34 and 38 by the experimental structure with PDB identifier (PDBid) 1A7W^30,33^ resolved at 1.55Å. None of those two were predicted as binding (at p≥0.5). Both the name and the available annotations suggested DNA-binding. If so, the observed metal-binding might point to allosteric binding. Four residues were predicted reliably (probability≥0.95) to bind nucleic acids (Fig. 5A, dark red residues). For another PDB structure of this protein (PDBid 5T5K^30,34^ at 4.0Å resolution), BioLiP annotates DNA-binding for all four reliably predicted residues. Due to our threshold in minimal resolution, this structure had not been included in our data sets. Overall, BioLiP annotates 13 residues in 5T5K as DNA-binding, 10 of those were correctly predicted (77% recall; Fig. 5A, lighter red). With respect to the three remaining: although our sequence-based method clearly did not reach remotely the power of X-ray crystallography, at least some of the parts of the proteins seemingly bridged over by the major grove (Fig. 5A: dark blue) might, indeed not bind DNA.

**Fig. 5:**
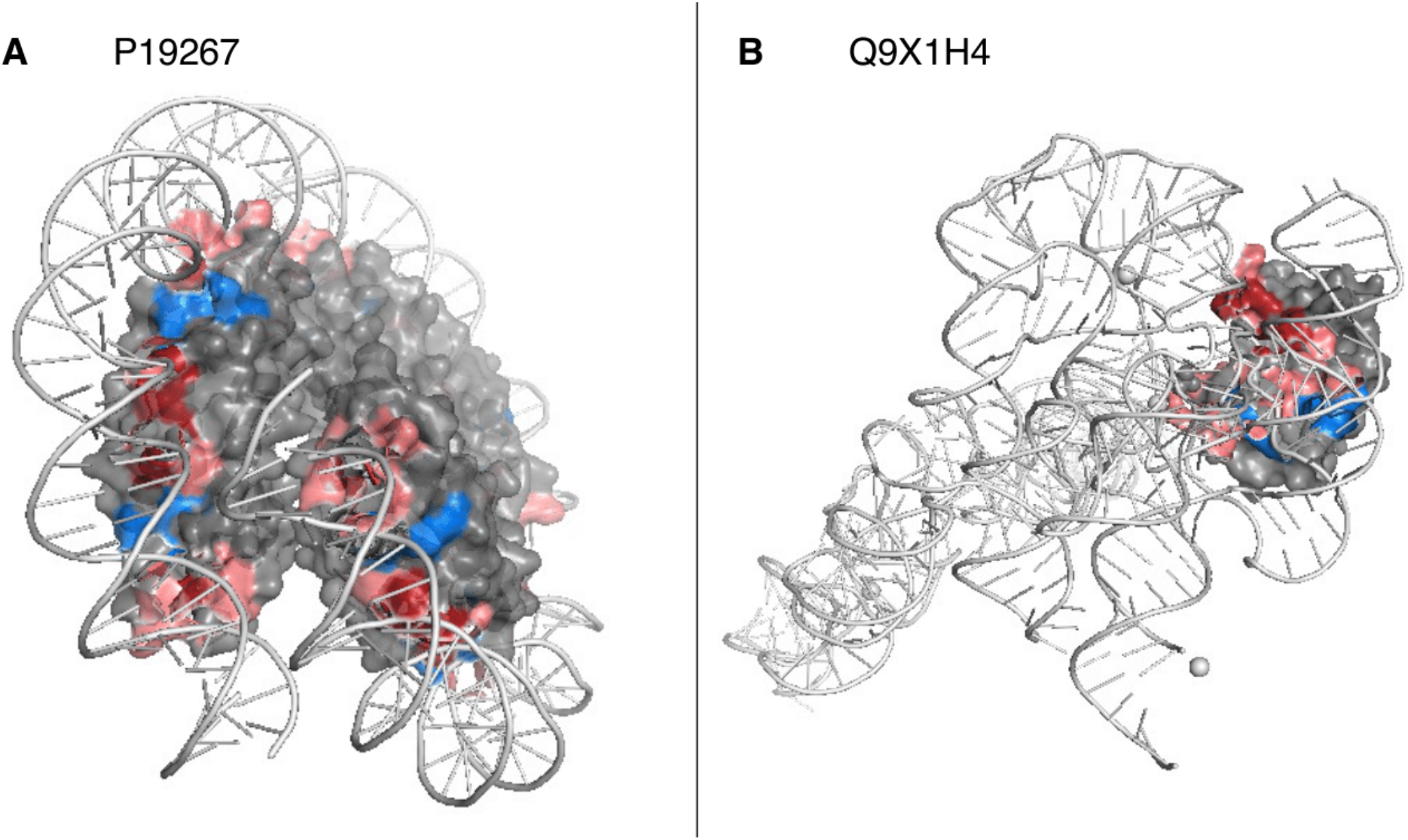
Annotations from low-resolution structures supported through reliable predictions. **A:** Our development set (DevSet1014) contained the PDB structure 1A7W^30,33^ for the DNA-binding protein HMf-2 (UniProt ID: P19267). No DNA/nucleic acid binding was annotated in that structure, but our new method, bindEmbed21DL, reliably predicted (probability ≥0.95) four residues to bind nucleic acids. Shown is the PDB structure 5T5K^30,34^ for the same protein that has a resolution of 4.0Å and annotations of DNA-binding, including the four most reliable predictions (dark red). Overall, 10 of 13 (77%) residues annotated as DNA-binding in 5T5K were also predicted by bindEmbed21DL (shown in lighter red; blue residues indicate experimental annotations which were not predicted). **B:** For the ribonuclease P protein component (UniProt ID: Q9X1H4), four residues were predicted with a probability ≥0.95 (indicated in dark red), none of these matched the annotations in the PDB structure 6MAX^30,35^. However, those four residues were considered as binding according to the two low-resolution structures 3Q1Q (3.8Å)^30,36^ (visualized) and 3Q1R (4.21Å)^30,36^. In total, those structures marked 21 binding residues; 15 of those 21 (71%) were correctly predicted (light red; blue residues observed to bind but not predicted). These two examples highlighted how combining low-resolution experimental data and very reliable predictions from bindEmbed21DL could refine those annotations and/or help designing new investigations.

We observed similar results for the ribonuclease P protein component (UniProt ID: Q9X1H4): the PDB structure 6MAX^30,35^ (1.42Å) annotated this protein with seven residues binding to a small molecule; none of the those were predicted at p≥0.95. Indeed, the available functional annotations clearly suggest nucleic acid-binding; the small molecule bound in 6MAX seems to mainly inhibit RNA-binding^35^. Four residues were predicted to bind nucleic acids reliably (p≥0.95, Fig. 5B, dark red). The low-resolution structures 3Q1Q (3.8Å)^30,36^ and 3Q1R (4.21Å)^30,36^ confirmed nucleic acid-binding for this protein. All four most reliable predictions were experimentally confirmed by those structures, and of the 21 residues annotated as binding, 16 were correctly predicted by default (p≥0.5, 76% recall, Fig. 5B, lighter red).

These two of 84 examples pitched *bindEmbed21DL* as a candidate tool to help in experimentally characterizing new binding residues completely different from the annotations it was trained on because all those correct binding predictions had been used as “non-binding” during training. On the one hand, this facilitates the identification of previously unknown binding sites; on the other hand, it might also help to verify and refine known, but potentially unreliable binding annotations, especially if multiple structures annotating different binding sites are available. In the two examples shown here, both proteins had already been annotated as binding to nucleic acids in less well-resolved structures, while the binding annotations from high-resolution structures rather pointed to binding of co-factors or inhibitors. Overall, the two examples suggested that the seemingly low performance values of *bindEmbed21DL* were, at least partially, rooted in the missing experimental annotations (residues not observed to bind treated as non-binding _DELETED_). We had selected the two of 84 by a simple algorithm: Pick those with an abundance of reliable binding predictions for which alternative experimental information was available. In doing this, we found that most seemingly incorrect binding predictions appeared correct. In fact, for those investigated in detail, precision was closer to 100% than to 80% (precision at p≥0.95, Fig. 4). Of the 84 proteins with seemingly incorrect, highly reliable binding predictions, 32 were predicted to bind nucleic acids. For 6 of those 32 proteins (19%), low resolution structures with binding annotations at least partially matching the predictions were available. On the other hand, only one of the 75 proteins with non-observed reliable (p≥0.95) metal predictions (1%) and one of the 80 proteins with non-observed reliable (p≥0.95) small molecule predictions were confirmed by low resolution structures. While those examples demonstrated qualitatively that our assessment clearly under-estimated performance, they did not suffice to adjust performance measures.

### Final method *bindEmbed21* combined HBI and ML

Homology-based inference (HBI) assumes that two sequence-similar proteins are evolutionary related, and therefore, also share a common function. Using HBI to predict binding residues for three different ligand classes for our validation set yielded very good results for low E-value thresholds, but at those thresholds, hits were only found for few proteins (Fig. S7). For instance, for E-values ≤ 10^−50^, HBI achieved F1=56±4% (Fig. S7, leftmost dark red bar), but only 198 of the 1,014 proteins found a hit, i.e., another protein with experimental annotations. When only using HBI to predict for all proteins, a random decision would have to be made for proteins without a hit. Thereby, performance dropped substantially (Fig. S7, leftmost light red bar). Hence, HBI outperformed our ML method *bindEmbed21DL* only for a small subset of proteins. We combined the best of both (*bindEmbed21DL* and HBI) applying a simple protocol: Predict binding residues through HBI if an experimentally annotated sequence-similar protein is available, otherwise use ML. This combination was best (highest recall) at an E-value threshold of 10^−3^ (Fig. S7A, blue bar). The optimum was not sharp; instead, numbers remained almost constant over at least six orders of magnitude in E-value. While this implied stability (other choices would have given similar results), one reason for the lack of a sharp optimum was the small data set combined with the fact that only about 5% of all residues bind. Therefore, increasing the E-value tenfold brought in much fewer binding residues than proteins (Fig. S8).

Combining ML and HBI improved performance on *TestSet300* by five percentage points for F1 (F1=48±3%; Fig. 6D, Table S8). HBI also improved performance for each ligand class (Fig. 6A-C, Table S8) except for the precision in predicting nucleic acid binding (Fig. 6B, Table S8). ML performance was somehow limited by overprediction, especially for metal ions and small molecules (low CovNoBind; Tables S3 & S7), i.e., many proteins were predicted to bind to those ligand classes without matching annotations. Combining bindEmbed21DL with HBI slightly reduced overprediction (higher CovNoBind, lower CovOneBind, Table S9) for all three ligand classes. Since the effect was largest for nucleic acids, this could explain the drop in precision of the final combined method *bindEmbed21* compared to the ML-only component *bindEmbed21DL*, because precision was set to zero for proteins annotated but not predicted to bind to a ligand class.

**Fig. 6:**
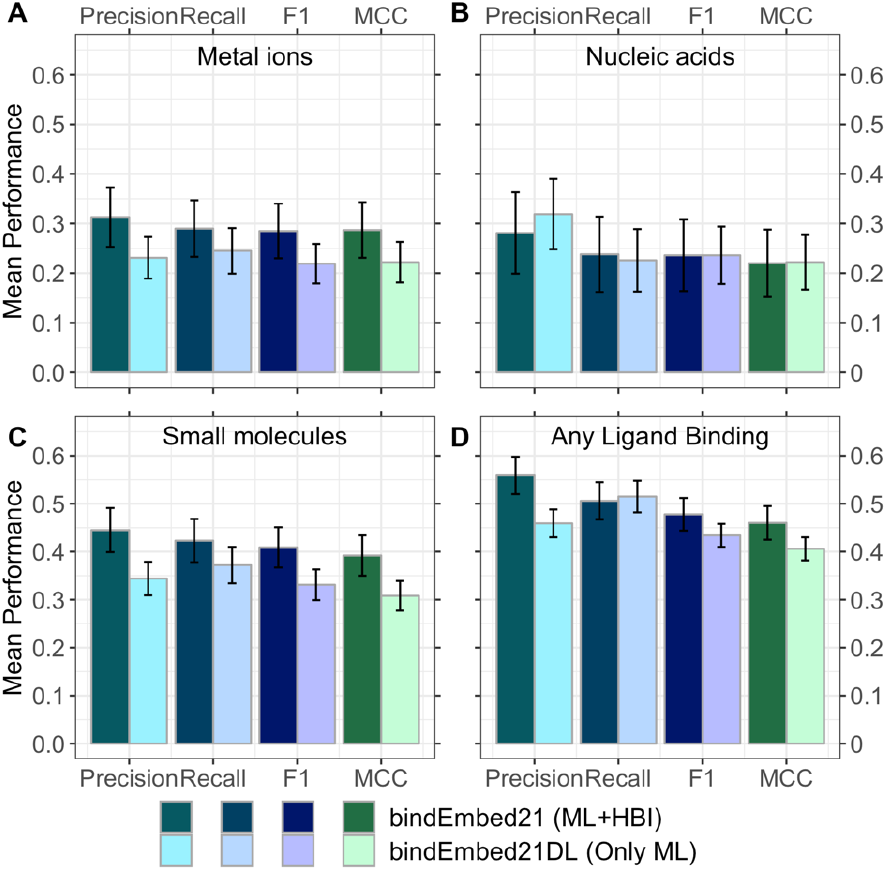
Best performance by combining ML and HBI. We combined homology-based inference (HBI) and Machine Learning (ML) by transferring annotations between homologs (E-value<10-3) if available and running de novo ML predictions using bindEmbed21DL, otherwise. This combination improved performance for the prediction of whether a residue binds to a certain ligand class for A. metal ions, B. nucleic acids, C. small molecules, and D. the combined, unspecific prediction of binding any of those three ligand classes vs. non-binding any of the three. The final version of bindEmbed21 achieved F1=29±6%, F1=24±7%, and F1=41±% for metal ions, nucleic acids, and small molecules, respectively. Lighter colored bars indicate the performance for the ML method, darker colors indicate the performance for the combination of ML and HBI.

### Prediction for complete human proteome discovered unknown candidate binding residues

Of the 20,386 sequences with 11,362,967 residues currently constituting the human proteome in Swiss-Prot^37^, only 3,121 (15%) had any structure with binding annotations in BioLiP (Fig. 7, Table S10). Using our protocol for HBI (transfer binding annotations of local alignment if E-value ≤ 10^−3^) transferred binding residues for another 7,199 proteins pushing the annotations from *BioLiP+HBI* to 51% (Fig. 7, Table S10; 53% for E-value cutoff of 1), i.e., for about half of all human proteins no ligand is known. As most proteins likely bind some ligand to function correctly, many ligands remain obscure. In fact, this calculation substantially under-estimated the extent of missing annotations by considering a single binding annotation as “protein covered” although 80% of the proteins have several domains^38,39^, i.e. there are not only missing annotations in the 49% of the proteins without annotation but also in other domains (or even other regions) of the proteins covered by *BioLiP+HBI*. Due to speed, applicability to three main ligand classes, and performance, *bindEmbed21DL* bridged this sequence-annotation gap predicting binding for 92% of the human proteins; for 42% of all human proteins (8,510), no binding information had been available without our prediction (Fig. 7, Table S10) and 21% of those 8,510 (1,751) were predicted reliably (probability≥0.95 corresponding to >73% precision, Table S7). In addition, for 21% of the proteins with experimental or HBI-inferred annotations, bindEmbed21DL provided highly reliable binding predictions previously unknown.

**Fig. 7:**
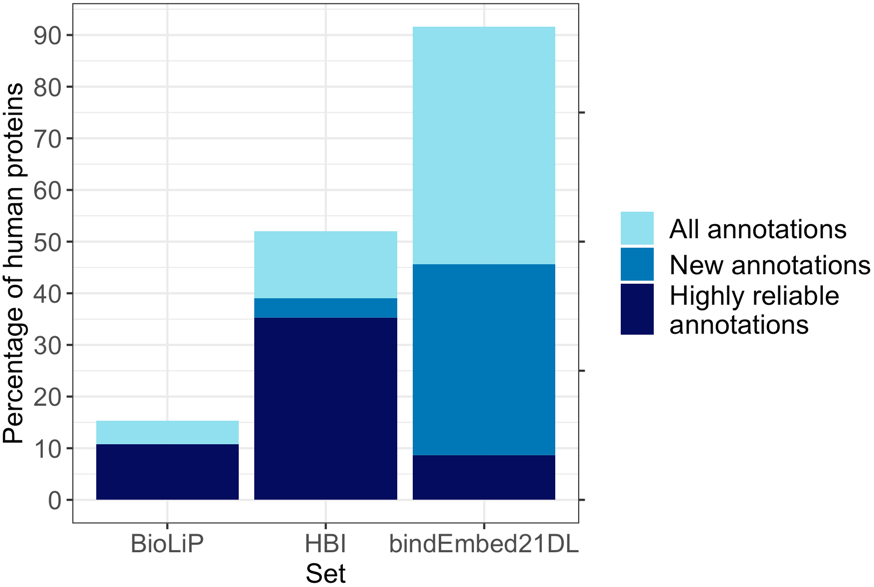
Binding predictions for complete human proteome. BioLiP: experimental annotations, HBI: homology-based inference, bindEmbed21DL: predictions. Data: human proteome from Swiss-Prot^37^ with 20,386 proteins. For 3,121 human proteins (15%), any binding annotation was experimentally known; 2,211 of those were reliable (resolution ≤2.5Å; darkblue bar for BioLiP). For 10,526 proteins (52%), binding annotations could be inferred using HBI at EVAL≤1 (light blue). Of those, 7,973 proteins were not previously annotated (blue bar); for 7,199 proteins without binding annotations, binding annotations could be inferred at EVAL≤10-3 following the protocol of bindEmbed21HBI (dark blue). Therefore, BioLiP+HBI allowed annotating some binding in 52% of the human proteome. bindEmbed21DL predicted binding residues for 18,663 proteins (92%) (light blue); no annotations were previously known (neither through BioLiP nor HBI) for 8,510 (blue). Highly reliable predictions (probability≥0.95) were possible for 1,751 proteins without previous binding annotations (dark blue).

Comparing the probability distributions of residues predicted to bind between proteins with and without annotations, we observed a clear difference between those (Fig. S9). Neither abundance in disordered regions nor abundance in membrane proteins nor different length distributions explained any aspect of the difference (Fig. S10). On the other hand, the large overlap between the distributions (Fig. S9) suggested that, while some of the newly predicted binding residues potentially stem from prediction mistakes, especially highly reliably predicted residues could point towards new binding residues.

One important result from the human proteome prediction was the relative contribution of the three ligand classes: Of all human residues, 1.2%, 2.0%, 3.1% were predicted to bind metal, nucleic acids, and small molecules, respectively (Tables S10&S11). Thus, about 20% of the binding residues were predicted to bind metal, 30% nucleic acid, and about 50% small molecules. Overall we assume that the mistakes made in all binding predictions were unbiased, i.e., the 20:30:50 (metal:nucleic:small) are likely good estimates for what a complete experimental coverage of all human proteins would reveal. This finding suggested that our *TestSet300* provided a much more representative mixture of these classes than *TestSetNew46* and a slightly more representative mixture than *DevSet1014* (Table S11).

As seen for the example of the human proteome, binding annotations are far from complete and cannot be inferred using HBI for most proteins leading to two major observations: (1) fast and generally applicable *de novo* prediction methods such as *bindEmbed21DL* are an important tool for the identification of new potential binding residues and ligands that could guide future experiments, and (2) our performance estimates are most likely too conservative due to missing annotations. In fact, while 48,700 residues were annotated as binding in structures with a resolution ≤2.5Å, an additional 21,057 residues were predicted as binding with a probability?0.95. Assuming that 15,372 of those are correct (precision at 0.95 is 73%, Table S7), our current set of annotations is likely missing 24% of binding residues.

Given its speed, bindEmbed21DL could also be easily applied to other complete proteomes. Predictions for all human proteins were completed within 80 minutes using one single Xeon machine with 400GB RAM, 20 cores and a Quadro RTX 8000 GPU with 48GB vRAM (40 minutes for the generation of the embeddings, 40 minutes for the predictions), i.e., generating binding residue predictions for one protein sequence took around 0.2 seconds allowing fast predictions for large sets of proteins.

### Availability

All data, the source code, and the trained model are available via GitHub (https://github.com/Rostlab/bindPredict). Embeddings can be generated using the bio_embeddings pipeline^40^. In addition, *bindEmbed21* and its components *bindEmbed21DL* and *bindEmbed21HBI* are publicly available through bio_embeddings. Users can apply the combined method or run its components independently. Therefore, binding residue predictions can be generated fully without the need of any alignment method.

## Conclusion

We proposed a new method, *bindEmbed21*, predicting whether a residue in a protein sequence binds to a metal ion, a nucleic acid (DNA or RNA), or a small molecule. The method combines homology-based inference (HBI: *bindEmbed21HBI*) with Artificial Intelligence (AI), in particular using input from deep learning (DL: *bindEmbed21DL*). *bindEmbed21DL* neither relied on knowledge of protein structure nor on expert-crafted features, nor on evolutionary information derived from multiple sequence alignments (MSAs). Instead, we inputted *embeddings* from the pre-trained protein Language Model (pLM) ProtT5^28^ into a two-layer CNN. The major problem with experimental data is the lack thereof: high-resolution data was available for fewer than 1,100 non-redundant proteins from any organism. Given the data sparsity, it is likely that many binding residues remain unknown even in the subset of 1,100 proteins with experimental data. Nevertheless, our evaluation equated “not observed” with “not binding”, treating predictions of non-observed binding as false positives. Although apparently blatantly underestimating precision, this crude simplification was needed to avoid over-prediction: methods only considering “what fraction of the experimental annotations is predicted?” (Recall, Eqn. 1) tend to optimize recall. The simplest non-sense path toward that end of “always predict binding” was carefully steered clear off by bindEmbed21DL which outperformed its MSA-based predecessor, bindPredictML17^5^, by 13 percentage points (Fig. 2A) and appeared competitive with the MSA-based method ProNA2020^19^ specialized to predict DNA- and RNA-binding and the zinc-binding prediction methods PredZinc^20^ and ZincBindPredict^29^ (Fig. 3). Prediction strength correlated with performance (Fig. 4): For the one fourth of all binding residues predicted with a probability≥0.95, 73% corresponded to experimentally known binding annotations available today (Table S7). Detailed analysis of very reliable predictions not matching known experimental annotations revealed that *bind*Embed21DL correctly predicted binding residues missing in the structures used for development (Fig. 5). The analysis of predictions for the entire human proteome underlined that most binding annotations remain unknown today (51% with binding annotations through experiments or homology) and that *bindEmbed21DL* can help in identifying new potential binding sites (Fig. 7, Table S10). Overall, about 6% of all residues in the human proteome were predicted to bind any of the three ligand classes covered (metals 1.2%, nucleotides 2.0%, small molecules 3.1%). The proteome analysis also suggested our performance estimates as too conservative: For the two carefully investigated case studies, many reliably predicted ligands not annotated tended to be correct. We combined the best from both worlds, namely AI/ML and HBI, to simplify predictions for users and to optimally decide when to use which (Fig. 6). The new method, *bindEmbed21*, is freely available, blazingly simple and fast, and apparently outperformed our estimates.

## Materials & Methods

### Data sets

Protein sequences with binding annotations were extracted from BioLiP^9^. BioLiP provides binding annotations for residues based on structural information from the Protein Data Bank (PDB)^30^, i.e., proteins for which several PDB structures with different identifiers exist may have multiple binding annotations. We extracted and combined (union) all binding information from BioLiP for all chains of PDB structures matching a given sequence, which have been determined through X-ray crystallography^41^ with a resolution of ≤2.5Å (≤0.25nm). All residues not annotated as binding were considered non-binding.

BioLiP distinguishes four different ligand classes: metal ions, nucleic acids (i.e., DNA and RNA), small ligands, and peptides. Here, we focused on the first three, i.e., on predicting the binding of metal ions, nucleic acids, or small ligands (excluding peptides). At point of accession (26-11-2019), BioLiP annotated 104,733 structures with high enough resolution and binding annotations which could be mapped to 14,894 sequences in UniProt^37^. This set was clustered to remove redundancy using UniqueProt^42^ with an HVAL<0 (corresponding to no pair of proteins in the data set having over 20% pairwise sequence identity over 250 aligned residues^43,44^). We provide more details on the data in Table S12 and on the redundancy reduction in Section 2.1 of the Supporting Online Material (**<u>SOM</u>**). The final set of 1,314 proteins was split into a development set with 1,014 proteins (called *<u>DevSet1014</u>* with 13,999 binding residues, 156,684 non-binding residues; Table S12) used for optimizing model weights and hyperparameters (after another random split into training and validation), and test set with 300 proteins (*<u>TestSet300</u>* with 5,869 binding residues, 56,820 non-binding residues; Table S12). To allow maximum overlap to the development set of bindPredictML17^5^, we first extracted the 225 proteins from the 1,314 proteins which were also part of the data set of bindPredictML17. Since bindPredictML17 was only trained on enzymes and DNA-binding proteins, this set was highly biased towards nucleic acid and small molecule binding (59 proteins binding to nucleic acids, 176 to small molecules, and 95 to metal ions). The additional 75 proteins were added to slightly adjust for this imbalance. However, a full adjustment was not possible without decreasing the size of the training set too much.

In addition, we created a new and independent test set by extracting all sequences with binding annotations which were added to BioLiP after our first data set had been built (deposited between 26 November 2019 and 03 August 2021). This yielded a promising 1,592 proteins. However, upon redundancy reduction with HVAL<0, this set melted down to 46 proteins with 575 binding and 5,652 non-binding residues (*<u>TestSetNew46</u>*; Table S12). These numbers imply two interesting findings: Firstly, about 17 experiments with binding data have been published every week over the last 91 weeks. Secondly, less than one experiment per week provided completely new insights into binding of residues not previously characterized in similar proteins (3% of all experiments). These observations underscored the importance of complementing experimental with *in silico* predictions.

### Protein representation and transfer learning

We used ProtT5-XL-UniRef50^28^ (in the following *ProtT5*) to create fixed-length vector representations for each residue in a protein sequence. The protein Language Model (pLM) ProtT5 was trained solely on unlabeled protein sequences from BFD (Big Fantastic Database; 2.5 billion sequences including meta-genomic sequences)^45^ and UniRef50^37^. ProtT5 has been built in analogy to the NLP (Natural Language Processing) T5^46^ ultimately learning some of the constraints of protein sequence. Features learned by the pLM can be transferred to any (prediction) task requiring numerical protein representations by extracting vector representations for single residues from the hidden states of the pLM (transfer learning). As ProtT5 was only trained on unlabeled protein sequences, there is no risk of information leakage or overfitting to a certain label during pre-training. To predict whether a residue is binding a ligand or not, we extracted 1024-dimensional vectors for each residue from the last hidden layer of ProtT5 (Fig. S13, Step 1) without fine-tuning it on the task of binding residue prediction (i.e., the gradient of the binding prediction was not backpropagated to ProtT5).

### AI/Deep Learning architecture

For *bindEmbed21DL*, we realized the supervised learning through a relatively shallow (few free parameters) two-layer Convolutional Neural Network (CNN; Fig. S13, Step 2). The CNN was implemented in PyTorch^47^ and trained with the following settings: Adamax optimizer, learning rate=0.01, early stopping, and a batch size of 406 (resulting in two batches). ProtT5 embeddings (from the last layer of ProtT5, 1024-dimensional vector per residue) were used as the only input. The first CNN layer consisted of 128 feature channels with a kernel (sliding window) size of k=5 mapping the input of size L x 1024 to an output of L x 128. The second layer created the final predictions by applying a CNN with k=5 and three feature channels resulting in an output of size L x 3, one channel per ligand class. A residue was considered as non-binding if all output probabilities were < 0.5. The two CNN layers were connected through an exponential linear unit (ELU)^48^ and a dropout layer^49^, with a dropout rate of 70%. The two-layer CNN proved to be the best-performing architecture among a variety of architectures including CNNs with more layers, feedforward neural networks, and combinations of both. Feature channels, learning rate, kernel size, and dropout rate were optimized using an exhaustive grid search.

To adjust for the substantial class imbalance between binding (8% of residues) and non-binding (92%), we weighted the cross-entropy loss function. Individual weights were assigned for each ligand class and were optimized to maximize performance in terms of F1 score (Eqn. 3) and MCC (Eqn. 4). Higher weights in the loss function increased recall (Eqn. 1), lower weights increased precision (Eqn. 2). The final weights were 8.9, 7.7, and 4.4 for metal ions, nucleic acids, and small molecules, respectively.

### Homology-based inference

Homology-based inference (HBI) generally proceeds as follows: Given a query protein Q of unknown binding and a protein E for which some binding residues are experimentally known, align Q and E; if the two have significant sequence similarity (SIM(Q,E)>T), transfer annotations from E to Q. The threshold T and the optimal way to measure the sequence similarity (SIM) are determined empirically. Most successful *in silico* predictions of function are predominantly based on HBI^4,8,50–55^.

In our case, we aligned query proteins without binding annotations with MMseqs2^56^, creating evolutionary profiles from the resulting multiple sequence alignments (MSAs) for each protein (family) (two MMseqs2 iterations, at E-value ≤ 10^−3^) against a 80% non-redundant database combining UniProt^37^ and PDB^30^ adapting a standard protocol based on PSI-BLAST^57^ which was implemented for other methods before^19,24,51^. The resulting profiles were then aligned at E-value ≤ 10^−3^ against a set of proteins with experimentally known binding annotations (see SOM 2.3 for explicit MMseqs2 commands). To save resources, we clustered the set of proteins with known annotations at 95% pairwise sequence identity (PIDE; PIDE(x,y)<95% for all protein pairs x, y). For performance estimates, self-hits were excluded. From all hits, the local alignment with the lowest E-value and highest PIDE to the query was chosen. If this hit contained any binding annotations in the aligned region, annotations were transferred between aligned positions, and all non-aligned positions in the query were considered as non-binding. If no binding annotations were located in the aligned region, the hit was discarded and no inference of binding annotations through homology was performed. Combining bindEmbed21HBI with the ML method bindEmbed21DL led to our final method, *bindEmbed21*.

### Performance evaluation

To assess whether a prediction was correct or not, we used the following standard annotations: True positives (TP) were residues correctly predicted as binding, false positives (FP) were incorrectly predicted as binding, true negatives (TN) were correctly predicted as non-binding, and false negatives (FN) were incorrectly predicted as non-binding. Based on this classification for each residue, we evaluated performance using recall (or sensitivity, Eqn. 1), precision (Eqn. 2), F1 score (Eqn. 3), and Matthews Correlation Coefficient (MCC, Eqn. 4).

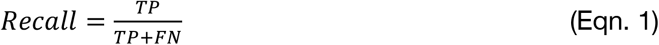

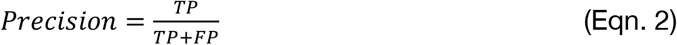

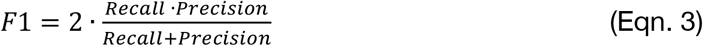

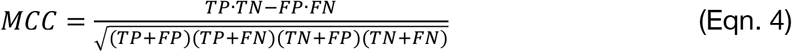

Negative recall (Eqn. 5), negative precision (Eqn. 6), and negative F1 score (Eqn. 7) focusing on the negative class, i.e., non-binding residues, were defined analogously:

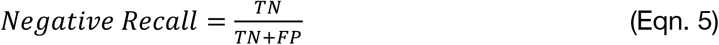

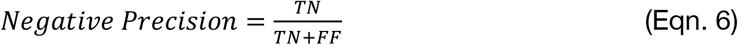

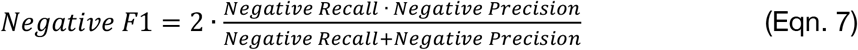

The measure *CovOneBind* (Eqn. 8) indicated the fraction of proteins for which at least one residue was predicted as binding. Accordingly, the inverse of this, *CovNoBind* (Eqn. 9), indicated the fraction of proteins for which predictions as well as experiments detected no binding. Since our data set only consisted of proteins with a binding site, *CovNoBind* had to be computed for different ligand classes, i.e., the fraction of proteins for which ligand *l* was neither observed nor predicted (Eqn. 9).

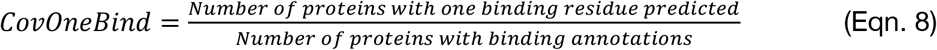

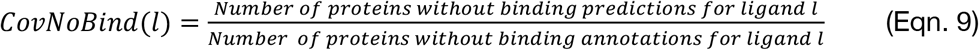

CovOneBind is an interesting measure to consider for experimentalists who submit only one sequence to a server and want to gauge how likely absence of prediction for that protein implies absence of binding. It does not give a clear indication of the performance of the method for a specific protein but attempts to capture how broadly applicable a method is. If a method only predicts binding residues for a small subset of proteins with high precision, it could still be considered inferior to a method predicting binding residues less precisely but for more proteins because those predictions can still provide valuable information.

Each performance measure was calculated for each protein individually. Then the mean was calculated over the resulting distribution and symmetric 95% confidence intervals (CI) assuming a normal distribution of the performance values were calculated as error estimates. While the performance values are not following a normal distribution, the sample size was sufficiently large to assume that a normal distribution can be applied in this case. For security we also tested bootstrapped CIs yielding the same results (SOM 2.4).

### Reliability Index

We transformed the probability *p* into a single-digit integer reliability index (RI) ranging from 0 (unreliable; probability=0.5) to 9 (very reliable; probability=1.0 for binding and probability=0.0 for non-binding) (Eqn. 10).

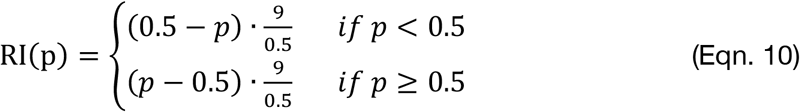

### Comparison to other methods

We compared our new method to the following four. We could not compare with other methods for different reasons (Table S14).

*<u>bindPredictML17</u>^5^* predicts binding residues from enzymes (trained on the PDB) and DNA-binding residues from PDIdb^31^. The method first builds MSAs and uses those to compute evolutionary couplings^23^ and effect predictions^21,22^. Those two main features are then used as input to the machine learning method.

*<u>ProNA2020</u>^19^* predicts binding to DNA, RNA, and other proteins using a two-step procedure: The first per-protein level predicts whether a protein binds DNA, RNA, or another protein. The second per-residue level predicts which residue binds to any (or all) of the three ligand classes. ProNA2020 combines HBI and machine learning using motif-based profile-kernel^58,59^ and word-based approaches (ProtVec)^60^ for the per-protein prediction and uses standard neural networks with different expert-crafted features taken from PredictProtein^24^ as input.

*<u>PredZinc</u>^20^* predicts binding to zinc ions using a combination of HBI inference and a Support Vector Machine (SVM). The SVM was trained on feature vectors representing the conservativity and physicochemical properties of single amino acids and pairs of amino acids.

<u>ZincBindPredict</u>^29^ is based on different Random Forest models to predict one particular zinc-binding site family. The models were trained on feature vectors encoding inter-residue distance, hydrophobicity, and number of charges around a residue.

## Abbreviations used

AI: artificial intelligence (expanding ML through deep learning, i.e., using more free parameters)
BFD: Big Fantastic Database (large database of protein sequences)
CI: confidence interval
CNN: Convolutional Neural Network
HBI: homology-based inference
(p)LM: (protein) language model
MCC: Matthews Correlation Coefficient
ML: machine learning
MSA: multiple sequence alignment
PDB: Protein Data Bank
PIDE: pairwise sequence identity
SOTA: state-of-the-art
SVM: support vector machine

## Acknowledgements

Thanks to Tim Karl and Inga Weise (both TUM) for invaluable help with technical and administrative aspects of this work. Last, but not least, thanks to all those who maintain public databases in particular Steven Burley (PDB, Rutgers), Ioannis Xenarios (Swiss-Prot, SIB, Geneva) and Yang Zhang (BioLiP, University of Michigan) and their crews, and to all experimentalists who enabled this analysis by making their data publicly available. This work was supported by the Bavarian Ministry of Education through funding to the TUM and by a grant from the Alexander von Humboldt foundation through the German Ministry for Research and Education (BMBF: Bundesministerium für Bildung und Forschung), by two grants from BMBF (031L0168 and program “Software Campus 2.0 (TUM) 2.0” 01IS17049) as well as by a grant from Deutsche Forschungsgemeinschaft (DFG-GZ: RO1320/4-1).

