## Supporting Online Material for "Protein embeddings and deep learning predict binding residues for various ligand classes"

### Table of Contents for Supporting Online Material

### Short description of Supporting Online Material

In this Supporting Online Material (SOM), we show a more thorough performance assessment and provide more details about the used data set and the underlying redundancy reduction (2.1), the Machine Learning (ML) method (Fig. S13), the MMseqs2 commands (Section 2.3), the calculation of error estimates (Section 2.4) and related work (Section 3).

Section 1.1 provides more details about the performance of bindEmbed21DL showing an assessment on various data sets (Table S1, Table S3), a comparison to random (Table S2) and a binarized version of bindEmbed21DL, namely bindEmbed21DL-binary (Table S6), and a more thorough analysis of the effect of over-prediction (Table S4) and cross-predictions (Table S5) on the overall performance of bindEmbed21DL.

Over-predictions could be reduced and therefore performance increased by only considering residues as binding if at least *x* residues were predicted as binding in this protein (Fig. S6). Additionally, the output probability of the method could be used to influence CovOneBind, CovNoBind, and precision (Section 1.5).

A more thorough comparison of different annotations used to define binding showed that the used annotations could highly affect the performance of a prediction method (Section 1.2&1.4, Fig. S3). However, the performance improvement of bindEmbed21DL over its predecessor bindPredictML17 was mainly due to replacing MSA-based input features with embeddings (Section 1.3, Fig. S2).

Combining bindEmbed21DL with homology-based inference (HBI) allowed an increase of precision and F1 even for high E-value thresholds, while recall dropped below the level of bindEmbed21DL for E-values >$10^{-3}$ (Fig. S7). Small changes in performance were due to only few new residues being inferred as binding for higher E-values (Fig. S8). Combining both approaches at an E-value cutoff of $10^{-3}$ led to an increase in CovNoBind but a drop in CovOneBind (Table S9).

bindEmbed21DL could be applied to obtain binding residues predictions for 92% of the human proteome (Section 1.7, Table S10, Table S11). A comparison of the distributions of prediction scores for experimentally verified binding residues, residues inferred through HBI, and previously unknown binding residues showed that previously unknown binding residues were predicted with on average slightly lower probability (Fig. S9). Neither an enrichment of disorder proteins nor transmembrane proteins nor a different length distribution could explain this difference (Fig. S10).

### 1. Additional Results

#### 1.1. Details on performance assessment of bindEmbed21DL.

**Table S1: Average performance for development set, test set, and new independent set. ***

| Set | | Precision | Recall | F1 | MCC |
|---|---|---|---|---|---|
| DevSet1014 | Metal ions | 25±3% | 27±3% | 24±2% | 0.24±0.02 |
| | Nucleic acids | 18±3% | 21±4% | 18±3% | 0.15±0.03 |
| | Small molecules | 27±2% | 30±2% | 26±2% | 0.23±0.02 |
| | Any ligand binding | 37±2% | 52±2% | 39±2% | 0.36±0.02 |
| TestSet300 | Metal ions | 23±4% | 25±5% | 22±4% | 0.22±0.04 |
| | Nucleic acids | 32±7% | 23±6% | 24±6% | 0.22±0.06 |
| | Small molecules | 34±4% | 37±4% | 33±3% | 0.31±0.03 |
| | Any ligand binding | 46±3% | 52±3% | 43±2% | 0.41±0.02 |
| TestSetNew46 | Metal ions | 25±14% | 28±15% | 26±14% | 0.3±0.1 |
| | Nucleic acids | 22±15% | 37±20% | 19±11% | 0.2±0.1 |
| | Small molecules | 31±10% | 34±12% | 29±9% | 0.25±0.09 |
| | Any ligand binding | 37±7% | 60±10% | 37±6% | 0.35±0.06 |

* We show precision, recall, F1, and MCC for the three sets used to evaluate bindEmbed21DL: Validation (DevSet1014), test (TestSet300), and a new independent test set of proteins added to BioLiP after November 2019 and non-redundant in itself and to the other two sets (TestSetNew46). Performance was similar for DevSet1014 and TestSetNew46, while bindEmbed21DL achieved better values for TestSet300. Therefore, bindEmbed21DL achieved good performance on the original data as well as succeeded in predicting binding residues for newer proteins. Error estimates indicate 95% confidence intervals.

**Table S2:Average performance for random approach on the test set. ***

| | | | | | |
|---|---|---|---|---|---|
| **TestSet300** | **Metal ions** | 2±1% | 1±1% | 1±1% | -0.01±0.01 |
| | **Nucleic acids** | 8±3% | 6±3% | 6±2% | 0.01±0.02 |
| | **Small molecules** | 7±1% | 7±1% | 6±1% | 0.00±0.01 |
| | **Any ligand binding** | 11±1% | 11±1% | 9±1% | 0.00±0.01 |

* We show precision, recall, F1 and MCC for the test set (TestSet300) using a random prediction. Random was generated by randomly shuffling the prediction probabilities of bindEmbed21DL. Error estimates indicate 95% confidence intervals.

Performance differed between ligand classes (Table S1). This could be due to differences in biophysical properties (i.e., small molecule binding was more clearly encoded in the embeddings) or due to differences in the data distribution (i.e., small molecule binding was more abundant in the development set, Table S12). To investigate, we re-trained bindEmbed21DL using a smaller development set of 515 proteins (DevSet515, Table S3) with only 108 proteins binding to small molecules. For this new set, performance of small molecule binding dropped immensely by 22 percentage points (Table S3). This suggested that rather data abundance than biophysical properties explained the difference in performance. If anything, it rather seems that nucleic acid binding was easier to predict due to the biophysical properties being more clearly encoded in the embeddings because this ligand class was predicted with an acceptable performance (Table S1) even though the number of proteins in DevSet1014 was fairly small compared to the other two classes (Table S12).

**Table S3: Average performance when training on subset of the development set. ***

| Set | | Precision | Recall | F1 | MCC |
|---|---|---|---|---|---|
| **DevSet515** | **Metal ions** | 33±3% | 39±4% | 34±3% | 0.34±0.03 |
| | **Nucleic acids** | 26±4% | 24±4% | 22±4% | 0.18±0.03 |
| | **Small molecules** | 12±4% | 3±1% | 4±2% | 0.05±0.02 |
| | **Any ligand binding** | 41±3% | 48±3% | 39±3% | 0.37±0.02 |

* We show precision, recall, F1, and MCC for a smaller development set (DevSet515) with only 108 proteins binding to small molecules. Training on this set, performance for small molecule binding dropped immensely indicating that this class was predicted better than the other classes on the original development

set (DevSet1014, Table S1) because it was overrepresented in the training set. Error estimates indicate 95% confidence intervals.

While for all three ligand classes for over 86% of the proteins at least one residue was predicted as binding (CovOneBind, Eqn. 8 in main text) (metal 86%, nucleic 93%, small 96%, Table S4), this high coverage of experimentally known ligands came from what appeared to be over-prediction as measured by the fraction of proteins not experimentally known (yet) to bind a particular ligand for which one was deemed to have been incorrectly predicted (1-CovNoBind(l), Eqn. 9 in main text): While binding to nucleic acids was only predicted for 19% of proteins without experimental data for nucleic acid binding (1-CovNoBind(nucleic acid)=100%-81%), this number rose to three fourth of the proteins for small molecules (Table S4). Metal ions and small molecules were also most often cross-predicted, i.e., residues in fact binding to small molecules were often predicted as binding to metal ions and vice versa (Table S5). This also explained the higher performance of the binary prediction (binding vs non-binding) compared to the performance for the individual ligand classes: Some residues were incorrectly predicted as binding to a certain ligand class and, therefore, were considered as false positives for this ligand class, but they could be in general involved in binding.

---

**Table S4: CovOneBind and CovNoBind for bindEmbed21DL. ***

| | **CovOneBind (Eqn. 8)** | **CovNoBind(l) (Eqn. 9)** |
|---|---|---|
| **Metal ions** | 86% | 37% |
| **Nucleic acids** | 93% | 81% |
| **Small molecules** | 96% | 25% |
| **Any ligand binding** | 99% | n/a |

* In each row, CovOneBind (Eqn. 8 in main text) indicates the number of proteins for which at least one residue was (correctly or incorrectly) predicted to bind to this ligand class (or any ligand class for the last row). CovNoBind(l) (Eqn. 9 in main text) is the percentage of proteins not annotated to bind to a certain ligand class for which also no residue was predicted as binding. Since the data set did not contain proteins without any binding annotations, the negative coverage is not defined for the general prediction of binding residues (last cell in the table). While bindEmbed21DL achieved a reasonable coverage, the negative coverage was low for metal ions and small molecules indicating that too many residues were predicted to bind to one of these two ligand classes. Data set: DevSet1014.

**Table S5: Confusion table of bindEmbed21DL for development set. ***

| | Metal ions | Nucleic acids | Small molecules | Non-Binding |
|---|---|---|---|---|
| **Metal ions** | **1,195 (34%)** | 56 (2%) | 598 (17%) | 1,670 (47%) |
| **Nucleic acids** | 67 (1%) | **1,647 (36%)** | 83 (2%) | 2,824 (61%) |
| **Small molecules** | 784 (6%) | 124 (1%) | **4,341 (33%)** | 7,725 (60%) |

* Rows indicate residues predicted by bindEmbed21DL as binding to a specific ligand; columns show the experimental (true) annotations. Values in the diagonal in bold font marked correct predictions. Most incorrect binding predictions were in fact non-binding residues. In addition, many residues predicted to bind metal ions are in fact binding to small molecules and vice versa. Data: DevSet1014.

Separately predicting whether a residue binds to a metal ion, a nucleic acid, or a small molecule is a more complicated prediction task than the binary distinction of binding and non-binding residues. To investigate whether performance could improve by only training on the binary task, we developed bindEmbed21DL-binary trained to distinguish binding from non-binding residues. On the same validation set as bindEmbed21DL, bindEmbed21DL-binary achieved F1=40±2%, i.e., one percentage point higher than bindEmbed21DL trained on three different ligand classes (Table S6). The two results could not be distinguished statistically, implying that the higher complexity in training on three ligand classes did not clearly affect performance. On the one hand, ML models tend to do better when applied to the same problem used for training, i.e., the class-agnostic method, *bindEmbed21DL-binary*, should have performed better. On the other hand, when the task is better defined, it is better to learn, i.e., the method trained on three classes, *bindEmbed21DL*, should have performed better. The observation of "no significant improvement" might have been the result of these two opposing trends.

**Table S6: Performance of bindEmbed21DL and bindEmbed21DL-binary. ***

| | Precision | Recall | F1 | MCC |
|---|---|---|---|---|
| **bindEmbed21DL** | 37±2% | 52±2% | 39±2% | 0.37±0.02 |
| **bindEmbed21DL-binary** | 37±2% | 57±2% | 40±2% | 0.36±0.02 |

* While being trained on the more complex task of distinguishing between three different ligand classes, bindEmbed21DL achieved F1=39±2% being only one percentage point worse than bindEmbed21DL-binary (F1=40±2%) which was only trained on predicting binding vs non-binding residues. All performance values are reported on the validation set. Error estimates indicate 95% confidence intervals.

##### 1.2. AI identified annotation errors.

Unlike bindEmbed21DL, bindPredictML17[1] was trained using annotations available through PDB[2] for enzymes and through PDIdb[3] for DNA-binding proteins. However, some binding annotations in the PDB might reflect crystal-induced rather than biologically relevant binding[4]. Therefore, we used annotations from BioLiP[4] for the training of bindEmbed21DL. Considering the predictions of bindPredictML17 for the 225 test proteins, we observed a better performance when using annotations from BioLiP for evaluation than when using annotations from PDB or PDIdb, although bindPredictML17 was trained on those annotations (Fig. 2A in the main text, lighter shaded bars higher than lightest shade bars). First, while training on noisy data, the seemingly false negative predictions of bindPredictML17 (Fig. 2B in the main text, rightmost bar labeled 'FN') were in fact often due to wrong annotations in the PDB. Without any re-training, the number of FN dropped by almost 40% when evaluating on annotations from BioLiP (Fig. 2B in the main text). Hence, bindPredictML17 had correctly captured incorrect binding annotations as non-binding. Secondly, these differences highlighted the importance of using high-quality binding annotations. Training on less noisy data might have been one reason for the improvement of bindEmbed21DL over bindPredictML17.

**Fig. S1: Seemingly false negative predictions in fact incorrect annotations.**

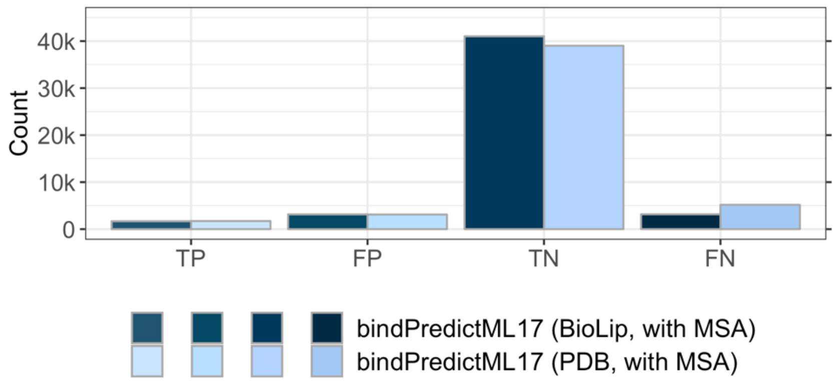

Investigating the number of true positives (TP), false positives (FP), true negatives (TN), and false negatives (FN) revealed that bindPredictML17 predicted many more FN when measured by PDB annotations than by BioLiP annotations. Hence, bindPredictML17 captured the incorrect binding annotations from the PDB correctly predicting those as non-binding which worsened its performance when assessing on those annotations but actually better captured the true binding residues. More details on the comparison of bindPredictML17 using BioLiP or PDB annotations can be found in SOM, Section 1.2.

##### 1.3. Performance gain mainly attributed to the replacement of MSAs with embeddings.

To investigate whether the performance gain of bindEmbed21DL over bindPredictML17 was mainly due to training on less noisy data or due to the replacement of MSA-based input features with embeddings, we re-trained bindEmbed21DL using the original training set of 412 proteins and the corresponding binding annotations of bindPredictML17. bindEmbed21DL-PDB already outperformed bindPredictML17 by, e.g., 13 percentage points in terms of F1 score (29±2% vs. 42±3%; Fig. S2). The replacement of PDB annotations with BioLiP annotations which also led to an increase in data set size from 412 to 1,014 resulted in a performance improvement of another five percentage points. Hence, training on BioLiP annotations instead of PDB annotations clearly improved performance, but the major gain in performance was achieved by replacing MSA-based features with data-driven inputs, namely embeddings.

---

**Fig. S2: Embeddings outperformed MSA-based predictions.**

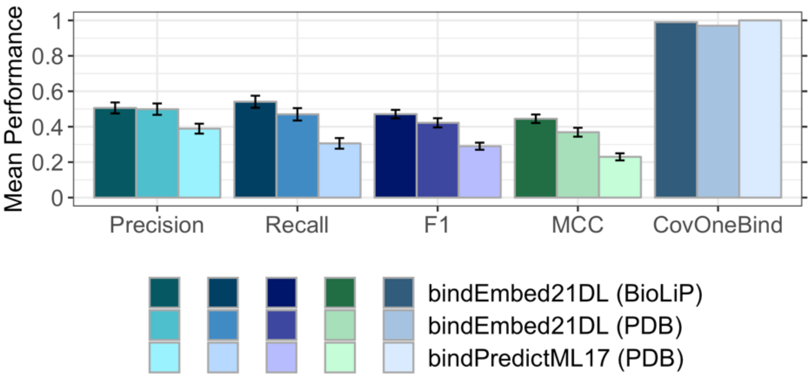

bindPredictML17 trained on a set of 412 proteins and PDB annotations achieved F1=29±2% (rightmost, lightest shaded bars). Training bindEmbed21DL on the same set but using embeddings as input improved performance by 13 percentage points leading to F1=42±3% (middle, darker shaded bars). Replacing PDB annotations with less noisy annotations from BioLiP improved performance by another five percentage points to F1=47±2% (leftmost, darkest shaded bars). This clearly showed that while using high-quality data was important, the major improvement was achieved by replacing MSA-based features with embeddings.

##### 1.4. Definition of binding highly influences performance.

In general, bindEmbed21DL achieved a higher F1 score and precision than ProNA2020[5], while ProNA2020 achieved a higher recall indicating that ProNA2020 predicted larger binding sites (see main text). ProNA2020 was trained on a different set of annotations obtained from PDIdb[3] and the Protein-RNA Interface Database (PRIDB)[6]. In this set, on average 21% of residues are annotated to bind to DNA or RNA compared to 12% for nucleic acid binding proteins in the test set of bindEmbed21DL. Therefore, ProNA2020 was trained on data where binding sites to DNA and RNA are more broadly defined, and therefore, consist of more binding residues leading to an over-prediction of binding residues from ProNA2020 for the test set of bindEmbed21DL. Since ProNA2020 was trained on different annotations, evaluating it using annotations from BioLiP is an unfair comparison. Therefore, we also assessed performance using the test set and annotations from ProNA2020. Using the 106 proteins binding to DNA or RNA from the test set of ProNA2020, ProNA2020 achieved F1=44±4% (Precision=45±5%, Recall=58±6%), while bindEmbed21DL-XNA achieved F1=38±5% (Precision=66±7%, Recall=32±5%) (Fig. S3). Therefore, bindEmbed21DL-XNA performed worse in terms of F1 score than ProNA2020 on its test set. However, the precision for bindEmbed21DL was significantly higher than for ProNA2020. Hence, the major difference between ProNA2020 and bindEmbed21DL seems to lie in the definition of what is involved in binding: While predictions from ProNA2020 focus on larger patches of binding residues, and therefore covering more of the actual binding site, bindEmbed21DL rather focuses on the prediction of key binding residues losing recall by making fewer predictions but resulting in more precise ones.

---

**Fig. S3: Comparison of bindEmbed21DL and ProNA2020 using binding annotations as defined by ProNA2020.**

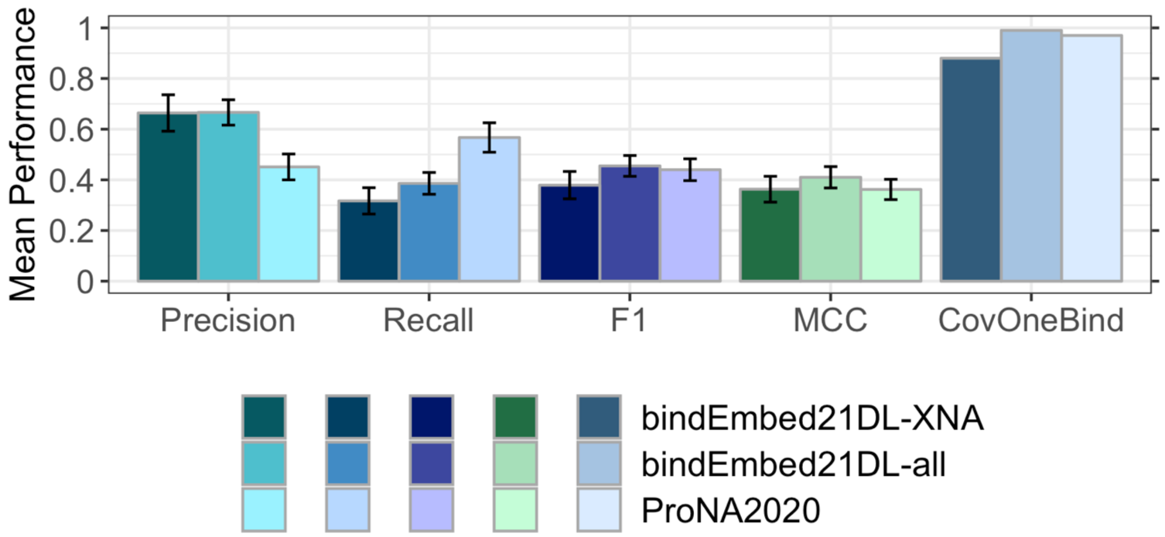

ProNA2020 (lightest shaded bars) was trained on a different set of annotations where, on average, 21% of residues were annotated to bind to DNA or RNA compared to 12% in the test set of bindEmbed21DL. To assess the effect of this different definition of binding, we evaluated performance using the test set and annotations from ProNA2020. Using the 106 proteins binding to DNA or RNA from the test set of ProNA2020, ProNA2020 achieved F1=44±4%, while bindEmbed21DL-XNA achieved F1=38±5%. Therefore, bindEmbed21DL-XNA performed worse than ProNA2020 in terms of F1, recall, and MCC on its test set.

However, the precision for bindEmbed21DL was significantly higher. Error bars indicate 95% confidence intervals.

##### 1.5. Refinement of predictions through focus on probability cutoff or number of predictions.

We analyzed the trade-off between precision, recall, and CovOneBind in dependence of the output probability of bindEmbed21DL for the different ligand classes. For higher cutoffs, precision increased, while CovOneBind dropped; the opposite trends were observed for lower cutoffs (Fig. S4). Based on the results for binding in general (Fig. 3 in the main text), we expected recall to increase for lower and decrease for higher cutoffs. However, the trend was not that consistent: While recall decreased as expected for higher cutoffs for small molecules (Fig. S4C), it first decreased and then increased for metal ions (Fig. S4A), and first increased and then decreased for nucleic acids (Fig. S4B). For proteins not binding to a certain ligand class *x* for which any residue was predicted to bind to *x,* precision and recall were set to 0. Increasing the cutoff to define a residue as binding decreased the number of residues incorrectly predicted to bind to *x.* Therefore, for more proteins not bound to *x,* there were also no residues predicted to bind to *x,* and those proteins were then ignored for the performance assessment (i.e., recall and precision are not set to 0). Therefore, recall could increase for higher cutoffs because CovNoBind increased (Fig. S4).

---

**Fig. S4: Performance of bindEmbed21DL for the three different ligand classes for different probability cutoffs.**

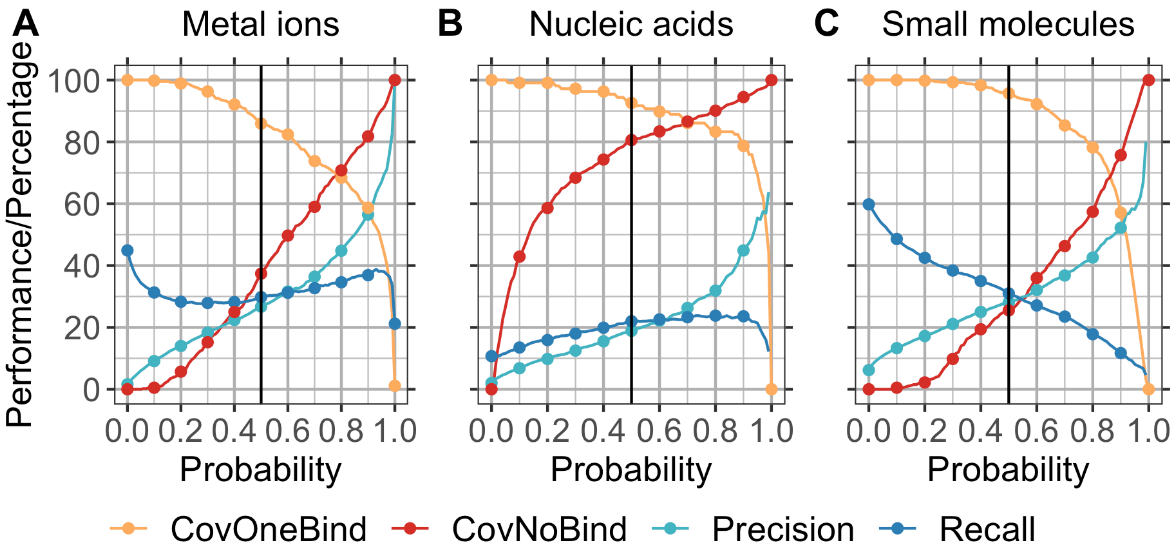

Residues were considered as binding to a certain ligand class if the output probability of bindEmbed21DL for this class was greater or equal to a specific cutoff. Choosing larger cutoffs led to an increase in precision and a decrease in coverage for **A.** metal ions, **B.** nucleic acids, and **C.** small molecules. The trend was not as clear for recall. While we would expect recall to decrease for higher cutoffs, it could also increase in this scenario due to an increase in negative coverage, i.e., if a residue is predicted to bind to a certain ligand class in a protein not binding to this class at all, recall is set to 0. If the number of such false positive predictions decreases (as it does for higher cutoffs), and therefore, less proteins are evaluated with a recall of 0, the recall could increase overall while actually decreasing for individual proteins. Black line at 0.5 marks performance for the default cutoff.

Seemingly incorrect binding predictions could in fact point towards new binding sites not yet experimentally verified. This is especially true for binding residues predicted with high probability ($p \geq 0.95$). To investigate whether this assumption holds, we compared 1024-dimensional ProtT5 embeddings and the internal representations from the first CNN layer (128 dimensions) of bindEmbed21DL for annotated binding residues, residues correctly predicted as binding (TP), and residues incorrectly predicted as binding (FP). The dimensionality of the input embeddings and representations from bindEmbed21DL was first reduced to 32 dimensions applying a Principle Component Analysis (PCA)[7] and was then further reduced to two dimensions using t-SNE[8]. For the original ProtT5 embeddings, falsely predicted binding residues formed wider spread clusters than correct predictions with the highly reliable predictions spread across those clusters for both false and correct binding predictions (Fig. S5B&C). Using the internal representations from bindEmbed21DL, clusters for false predictions were still more widely spread. However, highly reliable predictions were concentrated on the borders of the clusters (Fig. S5F). A similar pattern was observed for correct predictions with $p \geq 0.95$ (Fig. S5E). This indicated that highly reliable but false predictions were similar to correct predictions and could therefore point towards new potential binding residues.

**Fig. S5: t-SNE visualizations for ProtT5 embeddings and internal representations of the first CNN layer for binding annotations, true positive and false positive predictions.**

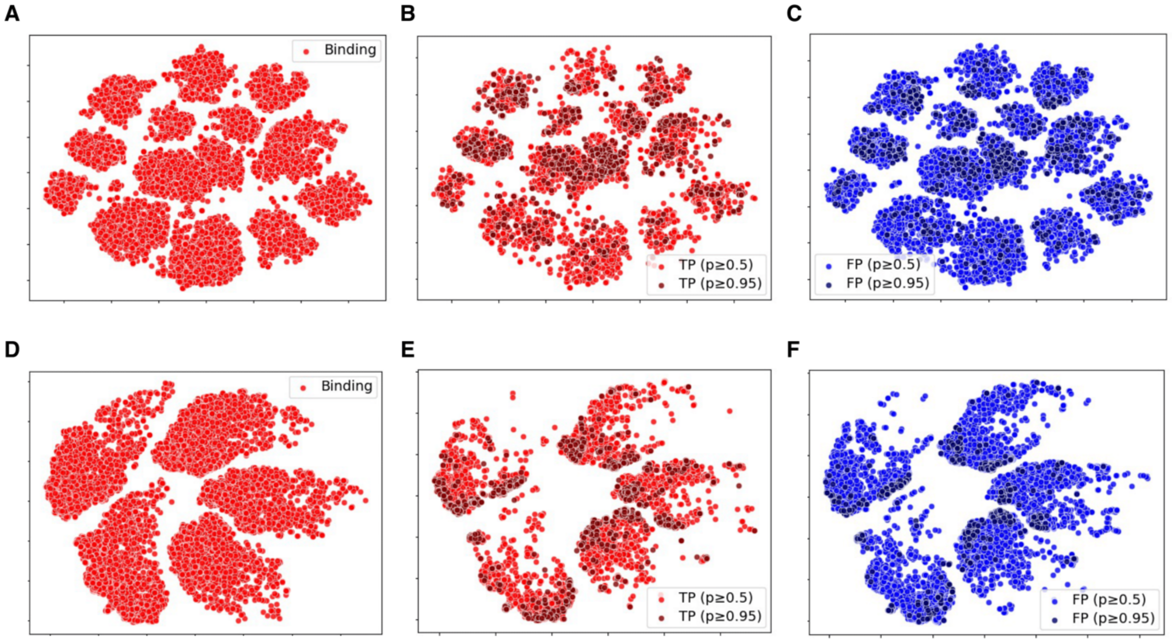

ProtT5 embeddings (1024 dimensions) and internal representations from the first CNN layer of bindEmbed21DL (128 dimensions) were first reduced to 32 dimensions using a PCA and were then further mapped to 2-dimensional representations using t-SNE. Those 2-dimensional representations were visualized for ProtT5 embeddings **(Panel A-C)** and representations from the first CNN layer **(Panel D-F)**. While all residues (including non-binding) were used to generate the 2-dimensional representations, we only visualize known binding residues **(Panel A and D)**, correctly predicted binding residues (TP; **Panel B and E**), and falsely correct binding residues (FP; **Panel C and F**). While highly reliable predictions were spread among all clusters for ProtT5 embeddings, they were more concentrated to the borders of the clusters for the internal representations of bindEmbed21DL. The similar patterns for highly reliable correct and false predictions indicated that highly reliable but incorrectly predicted binding residues could point towards new potential binding residues.

To provide binding predictions for as many proteins as possible, we considered a protein to bind to a specific ligand class if at least one residue was predicted to bind to this class. However, binding usually involves more than one residue, i.e., predicting only one residue as binding could indicate a wrong prediction. Predictions could be refined by only considering binding predictions if at least *x* residues were predicted to bind to this ligand class in a protein. Applying this filter led to an increase in CovNoBind(l) (Eqn. 9 in main text) for larger *x*, while decreasing CovOneBind (Eqn. 8; Fig. S6). While precision and recall were set to 0 for proteins annotated but not predicted to bind to a certain ligand class, those performance values still increased up to a certain threshold (Fig. S6; optimal threshold of 3, 10, and 8 residues for metal ions, nucleic acids, and small molecules, respectively). For those thresholds, more proteins falsely predicted to bind to this ligand class were removed than proteins actually binding to a certain ligand. Therefore, a low number of binding predictions in a protein indicated that those predictions were incorrect, and taking the number of predicted residues into consideration could help refining predictions (too few residues predicted: prediction less likely correct).

**Fig. S6: Performance of bindEmbed21DL in dependence of the minimum number of predictions considered.**

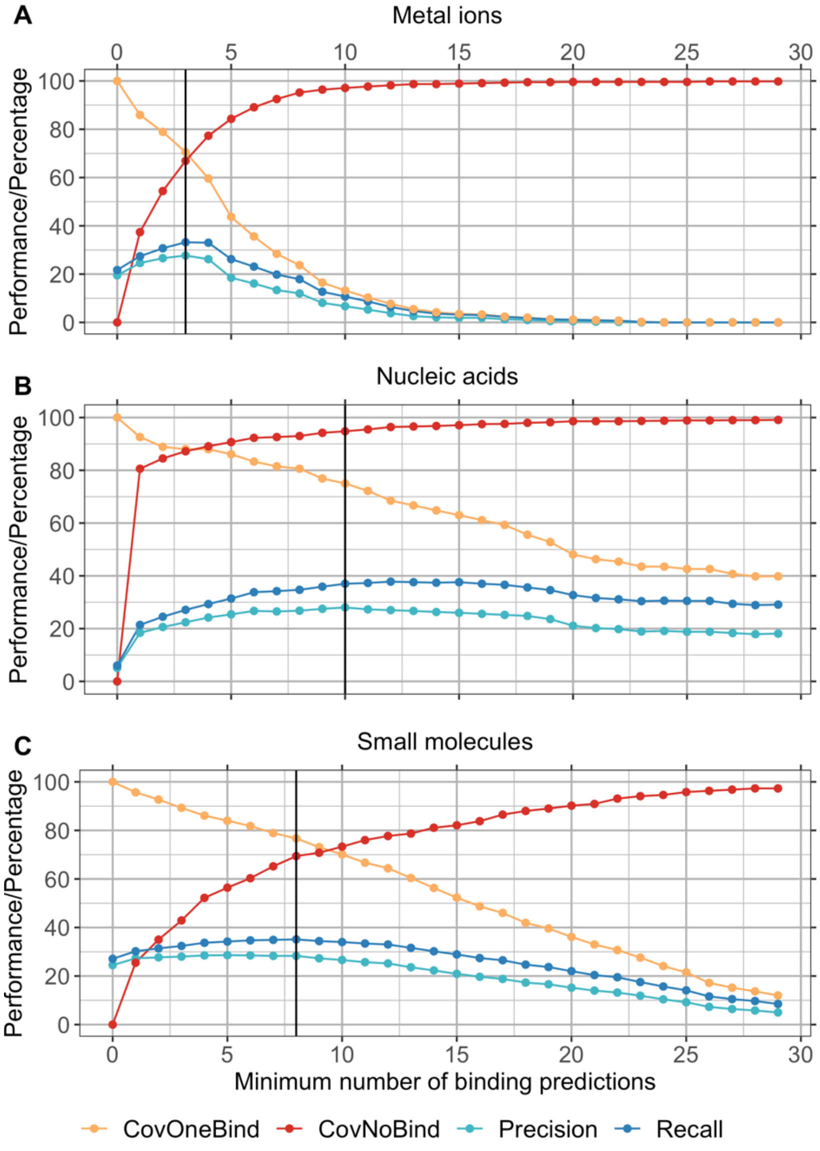

We show precision, recall, CovOneBind (Eqn. 8 in main text), and CovNoBind (Eqn. 9 in main text) if proteins were only considered to bind to a certain ligand class if at least *x* residues were predicted for this class, i.e., for proteins with <*x* binding predictions, we assumed that no binding residue was predicted. While no binding prediction was generated for more proteins (CovOneBind decreased) for larger *x,* CovNoBind increased because erroneous predictions were removed. Precision and recall also increased to a certain point (optimal x indicated by black, vertical line) indicating that proteins incorrectly predicted to bind to a ligand class had on average fewer binding predictions than proteins correctly predicted to bind.

#### Table S7: Performance of bindEmbed21DL for different probability cutoffs. *

| | Precision | Recall | F1 | Neg. Precision | Neg. Recall | Neg. F1 | CovOneBind |
|---|---|---|---|---|---|---|---|
| 0.0 | 10 | 100 | 16 | 0 | 0 | 0 | 100.0 |
| 0.02 | 12 | 96 | 20 | 95 | 29 | 42 | 100.0 |
| 0.04 | 14 | 93 | 23 | 96 | 42 | 57 | 100.0 |
| 0.06 | 16 | 90 | 25 | 97 | 51 | 65 | 100.0 |
| 0.08 | 17 | 87 | 27 | 97 | 57 | 70 | 100.0 |
| 0.1 | 19 | 84 | 28 | 97 | 54 | 68 | 100.0 |
| 0.12 | 20 | 82 | 30 | 97 | 65 | 77 | 100.0 |
| 0.14 | 21 | 80 | 31 | 97 | 69 | 79 | 100.0 |
| 0.16 | 22 | 78 | 32 | 97 | 72 | 81 | 100.0 |
| 0.18 | 23 | 76 | 33 | 97 | 74 | 83 | 100.0 |
| 0.2 | 24 | 74 | 33 | 97 | 76 | 84 | 100.0 |
| 0.22 | 25 | 72 | 34 | 96 | 78 | 85 | 100.0 |
| 0.24 | 26 | 71 | 35 | 96 | 79 | 86 | 100.0 |
| 0.26 | 27 | 69 | 35 | 96 | 81 | 87 | 100.0 |
| 0.28 | 28 | 67 | 36 | 95 | 82 | 88 | 100.0 |
| 0.3 | 29 | 66 | 37 | 96 | 83 | 88 | 99.9 |
| 0.32 | 30 | 64 | 37 | 96 | 84 | 89 | 99.9 |
| 0.34 | 31 | 63 | 37 | 96 | 85 | 90 | 99.8 |
| 0.36 | 32 | 62 | 38 | 96 | 86 | 90 | 99.7 |
| 0.38 | 33 | 61 | 38 | 95 | 87 | 91 | 99.7 |
| 0.4 | 34 | 59 | 39 | 95 | 88 | 91 | 99.5 |
| 0.42 | 34 | 58 | 39 | 95 | 89 | 91 | 99.4 |
| 0.44 | 35 | 57 | 39 | 95 | 89 | 92 | 99.2 |
| 0.46 | 36 | 56 | 39 | 95 | 90 | 92 | 99.1 |
| 0.48 | 37 | 55 | 39 | 95 | 91 | 92 | 98.8 |
| 0.5 | 38 | 53 | 39 | 95 | 91 | 93 | 98.6 |
| 0.52 | 39 | 52 | 39 | 95 | 92 | 93 | 98.3 |
| 0.54 | 39 | 50 | 39 | 95 | 92 | 93 | 98.2 |
| 0.56 | 40 | 49 | 39 | 94 | 93 | 93 | 98.0 |
| 0.58 | 41 | 48 | 39 | 94 | 93 | 93 | 97.9 |
| 0.6 | 42 | 47 | 39 | 94 | 94 | 94 | 97.4 |
| 0.62 | 43 | 46 | 39 | 94 | 94 | 94 | 97.0 |
| 0.64 | 45 | 45 | 39 | 94 | 94 | 94 | 96.3 |
| 0.66 | 46 | 44 | 39 | 94 | 95 | 94 | 95.6 |
| 0.68 | 47 | 42 | 39 | 94 | 95 | 94 | 95.2 |
| 0.7 | 48 | 41 | 39 | 94 | 95 | 94 | 94.0 |
| 0.72 | 49 | 40 | 39 | 93 | 96 | 94 | 92.6 |
| 0.74 | 50 | 39 | 38 | 93 | 96 | 94 | 91.0 |
| 0.76 | 51 | 38 | 38 | 93 | 97 | 95 | 90.0 |
| 0.78 | 53 | 37 | 38 | 93 | 97 | 95 | 88.9 |
| 0.8 | 54 | 35 | 37 | 93 | 97 | 95 | 87.3 |
| 0.82 | 57 | 34 | 37 | 93 | 97 | 95 | 85.2 |
| 0.84 | 59 | 33 | 36 | 93 | 98 | 95 | 82.8 |
| 0.86 | 60 | 31 | 36 | 92 | 98 | 95 | 80.0 |
| 0.88 | 62 | 30 | 35 | 92 | 98 | 95 | 75.9 |
| 0.9 | 65 | 29 | 34 | 92 | 98 | 95 | 71.2 |
| 0.92 | 69 | 27 | 33 | 92 | 99 | 95 | 65.6 |
| 0.94 | 72 | 26 | 33 | 92 | 99 | 95 | 56.6 |
| 0.95 | 73 | 25 | 32 | 92 | 99 | 95 | 51.2 |
| 0.96 | 75 | 25 | 33 | 92 | 99 | 95 | 44.9 |
| 0.97 | 76 | 25 | 33 | 91 | 99 | 95 | 37.9 |
| 0.98 | 78 | 25 | 33 | 91 | 99 | 94 | 29.3 |
| 0.99 | 81 | 23 | 32 | 90 | 99 | 94 | 18.9 |

* We show performance values for (negative) precision, (negative) recall, (negative) F1 score and CovOneBind for different probability cutoffs. Values marked in orange are discussed in the main text, value marked in dark orange corresponds to the default cutoff of 0.5. Values marked in grey indicate probability steps of 0.1 for easier readability.

##### 1.6. Combination of bindEmbed21DL with homology-based inference.

**Fig. S7: Performance of homology-based inference for different E-value thresholds.**

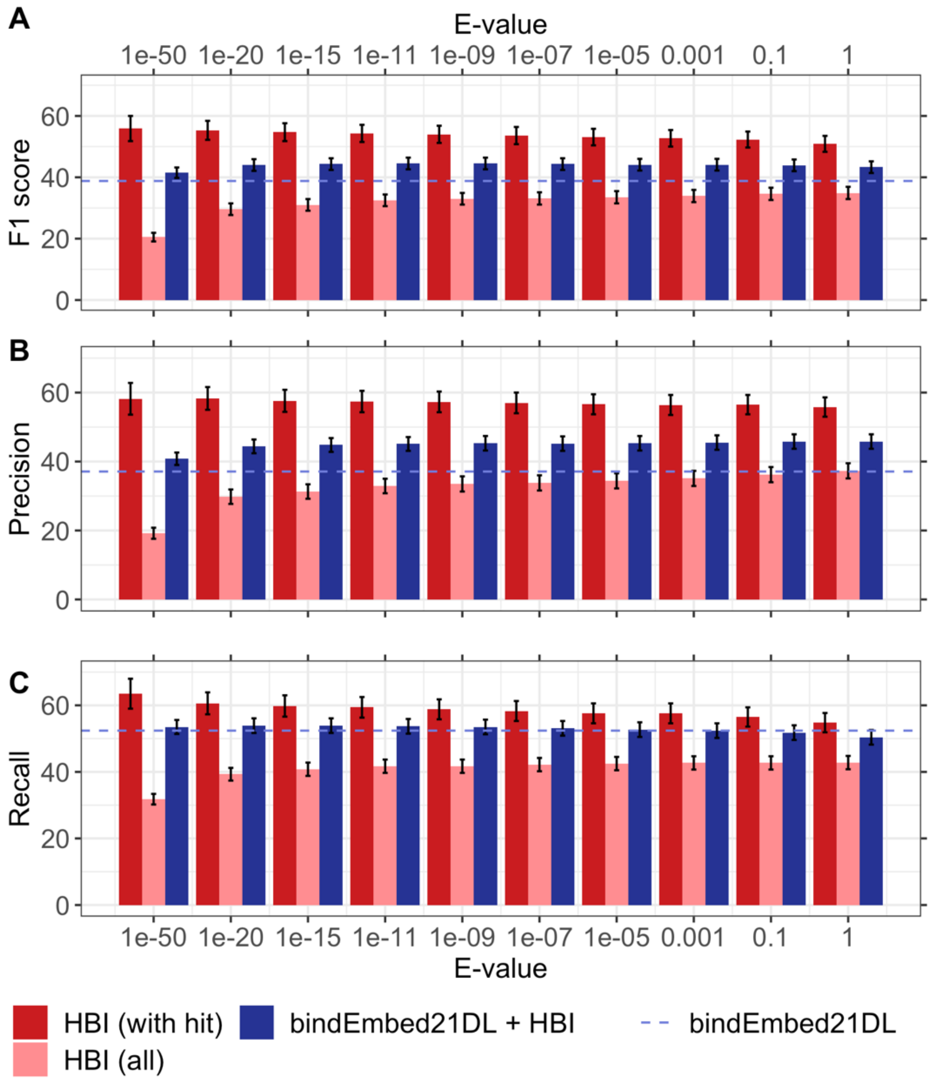

Performance for homology-based inference (HBI) as measured by **A.** the F1 score, **B.** the precision, and **C.** the recall varied with the E-value thresholds (red bars). The highest F1 of 56±4% was reached at E-value ≤ $10^{-50}$. However, if forcing predictions for all proteins by assigning binding residues at random if no homolog was available, F1 dropped to 21±2%

(leftmost light red bar). The combination of HBI with bindEmbed21DL (blue bars) performed numerically best for E-value $\leq 10^{-9}$ achieving F1=45±2%. However, performance values behaved similarly for all three measures (F1, precision, recall). To allow annotation transfer for the largest number of proteins possible without having the performance drop below that of bindEmbed21DL, we chose a final E-value threshold of $10^{-3}$ where F1 and precision are higher than for bindEmbed21DL (dashed line) and the recall is the same. Error bars indicate 95% confidence intervals.

---

**Fig. S8: Number of proteins and number of binding residues inferred through homology-based inference.**

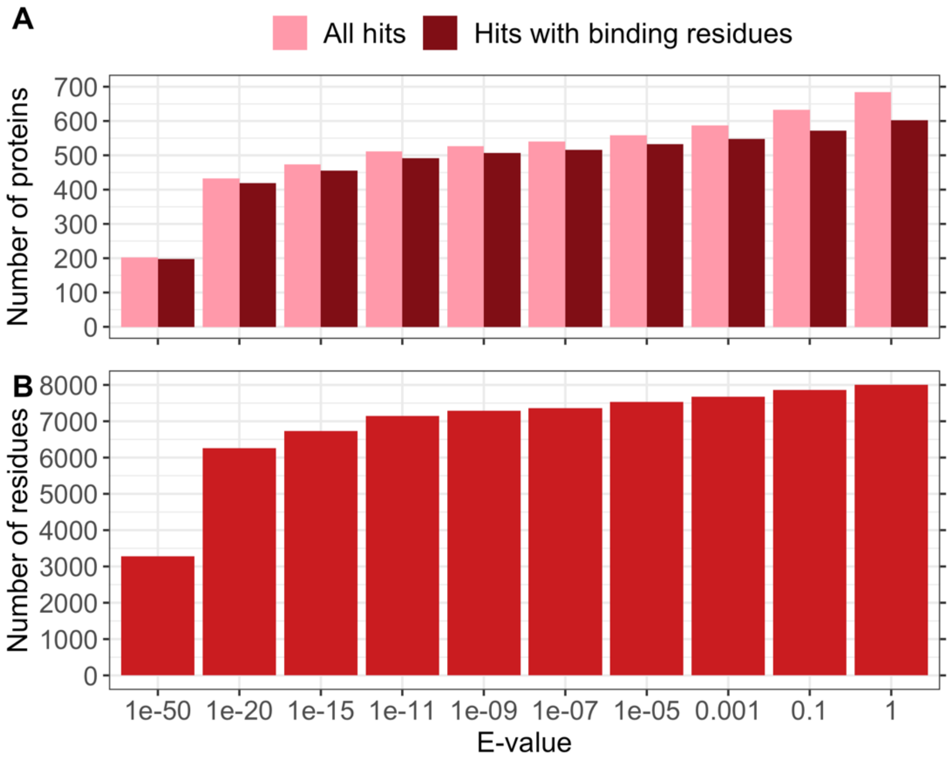

**A.** For lower E-value thresholds, binding residues could be inferred through homology-based inference (HBI) for fewer proteins. With increasing E-values, the number of hits increased. However, for some proteins, the local alignment did not contain any binding annotations, and those hits were discarded (difference between light and darker red). **B.** For many higher E-values, the increase in the number of inferred binding residues was small. This also explained why we did not observe a difference in performance for these different E-values (Fig. S7).

**Table S8: Average performance for bindEmbed21DL and bindEmbed21. ***

| | | Precision | Recall | F1 | MCC |
|---|---|---|---|---|---|
| bindEmbed21DL | Metal ions | 23±4% | 25±5% | 22±4% | 0.22±0.04 |
| | Nucleic acids | 32±7% | 23±6% | 24±6% | 0.22±0.06 |
| | Small molecules | 34±4% | 37±4% | 33±3% | 0.31±0.03 |
| | Any ligand binding | 46±3% | 52±3% | 43±2% | 0.41±0.02 |
| bindEmbed21 | Metal ions | 31±6% | 29±6% | 29±6% | 0.29±0.06 |
| | Nucleic acids | 28±8% | 24±8% | 24±7% | 0.22±0.07 |
| | Small molecules | 45±5% | 42±5% | 41±4% | 0.39±0.04 |
| | Any ligand binding | 56±4% | 51±4% | 48±3% | 0.46±0.04 |

* We show precision, recall, F1 and MCC for bindEmbed21DL (only using the Machine Learning (ML) method) and bindEmbed21 (combining ML with homology-based inference). Error estimates indicate 95% confidence intervals. Data set: TestSet300.

**Table S9: CovOneBind and CovNoBind for bindEmbed21DL and bindEmbed21. ***

| | bindEmbed21DL (only ML) | | bindEmbed21 (ML+HBI) | |
|---|---|---|---|---|
| | CovOneBind (Eqn. 8) | CovNoBind(l) (Eqn. 9) | CovOneBind (Eqn. 8) | CovNoBind(l) (Eqn. 9) |
| Metal ions | 96% | 30% | 82% | 62% |
| Nucleic acids | 77% | 89% | 53% | 95% |
| Small molecules | 94% | 20% | 87% | 55% |
| Any ligand binding | 98% | n/a | 99% | n/a |

* In each row, CovOneBind (Eqn. 8 in main text) indicates the number of proteins for which at least one residue was (correctly or incorrectly) predicted to bind to this ligand class (or any ligand class for the last row). The CovNoBind(l) (Eqn. 9 in main text) is the percentage of proteins not annotated to bind to a certain ligand class for which also no residue was predicted as binding. Combining bindEmbed21DL with HBI led to an increase in CovNoBind(l) but a drop in CovOneBind. Since HBI only used binding annotations from one local alignment, binding to multiple ligand classes is hard to predict because we could not identify different binding sites not close in the sequence. Data set: TestSet300.

#### 1.7. Full proteome prediction allows identification of previously unknown binding residues.

**Table S10: Binding information for human proteome. ***

| | | Overall | Metal ions | Nucleic acids | Small molecules |
|---|---|---|---|---|---|
| **Annotations (BioLiP, $\leq 2.5$Å)** | # Proteins with binding | 2,211 (11%) | 1,130 | 231 | 1,640 |
| | % of binding residues per protein | 5.5% (0.4%) | 2.0% | 5.4% | 5.7% |
| **Annotations (BioLiP, all structures)** | # Proteins with binding | 3,121 (15%) | 1,618 | 506 | 2,089 |
| | % of binding residues per protein | 5.7% (0.6%) | 2.1% | 6.4% | 5.6% |
| **HBI (E-value $\leq 10^{-3}$)** | # Proteins with binding | 9,694 (48%) | 4,746 | 1,622 | 6,365 |
| | # New proteins with binding | 7,199 (35%) | 3,763 | 1,381 | 4,811 |
| | % of binding residues per protein | 4.4% (1.6%) | 1.8% | 4.8% | 4.4% |
| **HBI (E-value $\leq 1$)** | # Proteins with binding | 10,526 (52%) | 5,121 | 1,695 | 7,018 |
| | # New proteins with binding | 7,973 (39%) | 4,127 | 1,448 | 5,436 |
| | % of binding residues per protein | 4.4% (1.7%) | 1.7% | 4.7% | 4.5% |
| **All annotations + HBI (E-value $\leq 10^{-3}$)** | # Proteins with binding | 10,320 (51%) | 5,381 | 1,887 | 6,900 |
| | % of binding residues per protein | 5.1% (1.9%) | 2.0% | 5.4% | 5.0% |
| **Predictions using bindEmbed21DL** | # Proteins with binding | 18,663 (92%) | 14,301 | 6,190 | 14,411 |
| | # New proteins with binding | 8,510 (42%) | 9,419 | 4,567 | 8,063 |
| | % of binding residues per protein | 3.9% (3.1%) | 1.2% | 2.0% | 3.1% |
| **Highly reliable predictions from bindEmbed21DL** | # Proteins with binding | 5,962 (29%) | 3,698 | 1,310 | 1,529 |
| | # New proteins with binding | 1,751 (9%) | 1,503 | 556 | 520 |
| | % of binding residues per protein | 1.0% (0.2%) | 0.7% | 1.0% | 0.7% |

* We show the number of proteins from the 20,386 sequences in the human proteome with binding information and the percentage of binding residues of (i) proteins with binding information and (ii) all proteins (in brackets). Using all available information from BioLiP (2nd row), 15% could be annotated with binding

information. Homology-based inference (HBI) (3rd row) adds another 36%. bindEmbed21DL provides predictions for another 42% corresponding to 8,510 proteins (5th row). Of those 8,510 proteins, 5,962 proteins contain highly reliable binding predictions (residues predicted with a probability ≥ 0.95), i.e., for 29% of the human proteome, highly reliable binding predictions could be provided by bindEmbed21DL while no annotations were available from experiments or homologs.

**Table S11: Percentage of predicted residues for human, DevSet1014, TestSet300, and TestSetNew46. ***

| | | **Metal ions** | **Nucleic acids** | **Small molecules** | **Total** |
|---|---|---|---|---|---|
| **Human** | *Predicted residues* | 1.2% | 2.0% | 3.1% | 6.3% |
| | *% of 3 classes* | 19.0% | 31.7% | 49.2% | 100% |
| | *Reliably predicted residues* | 0.7% | 1.0% | 0.7% | 2.4% |
| | *% of 3 classes* | 29.2% | 41.7% | 29.2% | 100% |
| **DevSet1014** | *Predicted residues* | 2.1% | 2.7% | 7.5% | 12.3% |
| | *% of 3 classes* | 16.6% | 18.7% | 64.7% | 100% |
| **TestSet300** | *Predicted residues* | 1.6% | 1.4% | 6.6% | 9.6% |
| | *% of 3 classes* | 14.1% | 23.5% | 62.4% | 100% |
| **TestSetNew46** | *Predicted residues* | 2.0% | 5.6% | 7.2% | 14.8% |
| | *% of 3 classes* | 13.3% | 13.3% | 73.4% | 100% |

* We show the percentage of predicted residues in each ligand class (metal ions, nucleic acids, small molecules) and the percentage of all three classes those residues account for (predicted residues/total). The composition for the human proteome (20:30:50 for metal:nucleic:small) was most similar to TestSet300. **Data sets:** Human: Predictions for 92% of the human proteome (Table S11); DevSet1014: Development set with 1,014 proteins, TestSet300: Test set with 300 proteins; TestSetNew46: New independent set with 46 proteins.

For almost half of the human proteins, no binding annotation is known, and for previously annotated proteins, many residues were newly predicted as binding. This could indicate a high prediction error of our method. On the other hand, those predictions could indicate previously unknown binding sites. The distributions of prediction scores for residues predicted as binding and (i) annotated as binding or (ii) inferred as binding through homology-based inference were similar (Fig. S9), while residues not annotated or inferred as binding were on average predicted with lower scores (Fig. S9). We expected a certain shift to the left (i.e., to 0.5) because the residues predicted as binding without any annotations will contain some false positive predictions which are predicted less reliably (Fig. 4 in the main text). Also, proteins with known or inferred binding annotations could have been similar to proteins in our training set. However, also other aspects could lead to this shift in the observed prediction probabilities. We investigated whether predictions were more difficult for (i) disordered proteins, (ii) transmembrane proteins, or (iii) proteins with different length than in the development set. Disorder was calculated using MetaDisorder[9], and we removed all proteins with at least 30 consecutive disordered residues to obtain a set of ordered proteins. Transmembrane helices (TMH) were predicted using TMSEG[10], and we excluded every protein with at least two TMHs to obtain a set of non-membrane proteins. In addition, for each of the three sets (proteins with experimentally verified annotations, proteins with inferred annotations, proteins with no annotations), we drew a subset of proteins mirroring the length distribution of the development set. None of these three aspects could explain the observed shift in distributions (Fig. S10). While this analysis did not reveal any insights whether new binding predictions could indicate previously unknown binding residues, it clearly showed that our method was not biased to ordered proteins, non-membrane proteins, or proteins of a specific length. Since no bias in the data set explained the shift in distribution, some of the shift is most likely explained by prediction mistakes, i.e., the large fraction of residues predicted with a probability close to 0.5 is probably pointing to wrong predictions. On the other hand, the distributions overlap to a certain extent, and especially residues predicted with a large probability could still point towards previously unknown binding residues.

**Fig. S9: Distribution of prediction scores for predicted binding residues annotated and not annotated as binding.**

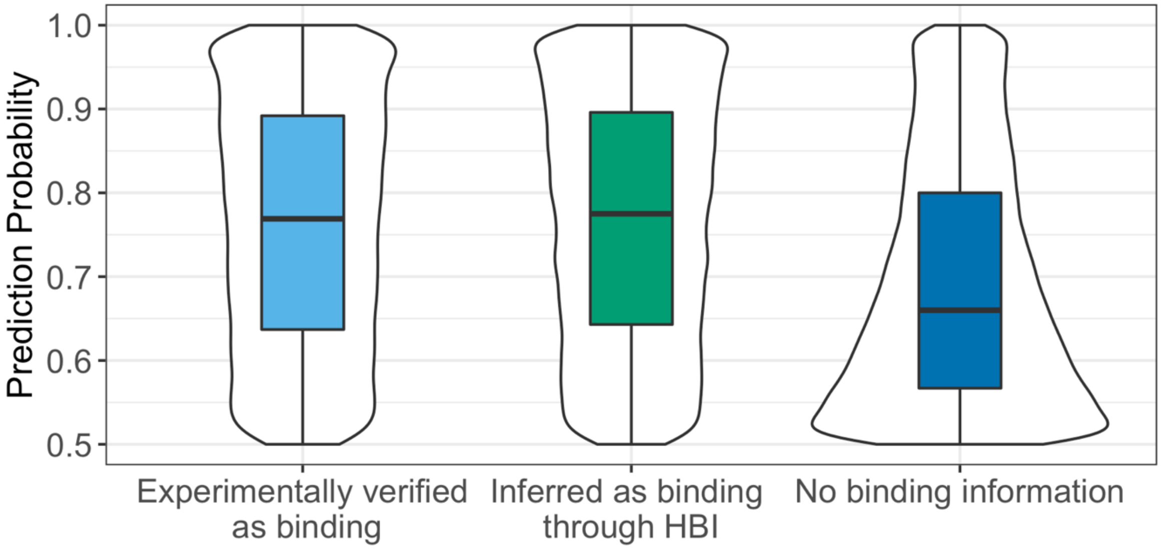

Residues which were not experimentally verified as binding residues or could be inferred through homology-based inference (HBI) were on average predicted with lower probability as binding (dark blue box lower than the other 2 boxes). Proteins with any binding information (either annotated or inferred) were similar to proteins in our training set. Therefore, bindEmbed21DL had seen those data points before and could make more reliable predictions. However, also some residues without any binding annotation could be predicted reliable and the distributions for all three sets overlapped to a large extent indicating that those residues not annotated as binding but predicted as such did not only originate from prediction mistakes but could indicate previously unknown binding annotations.

**Fig. S10: Distribution of prediction scores for ordered proteins, non-membrane proteins, and proteins of specific length.**

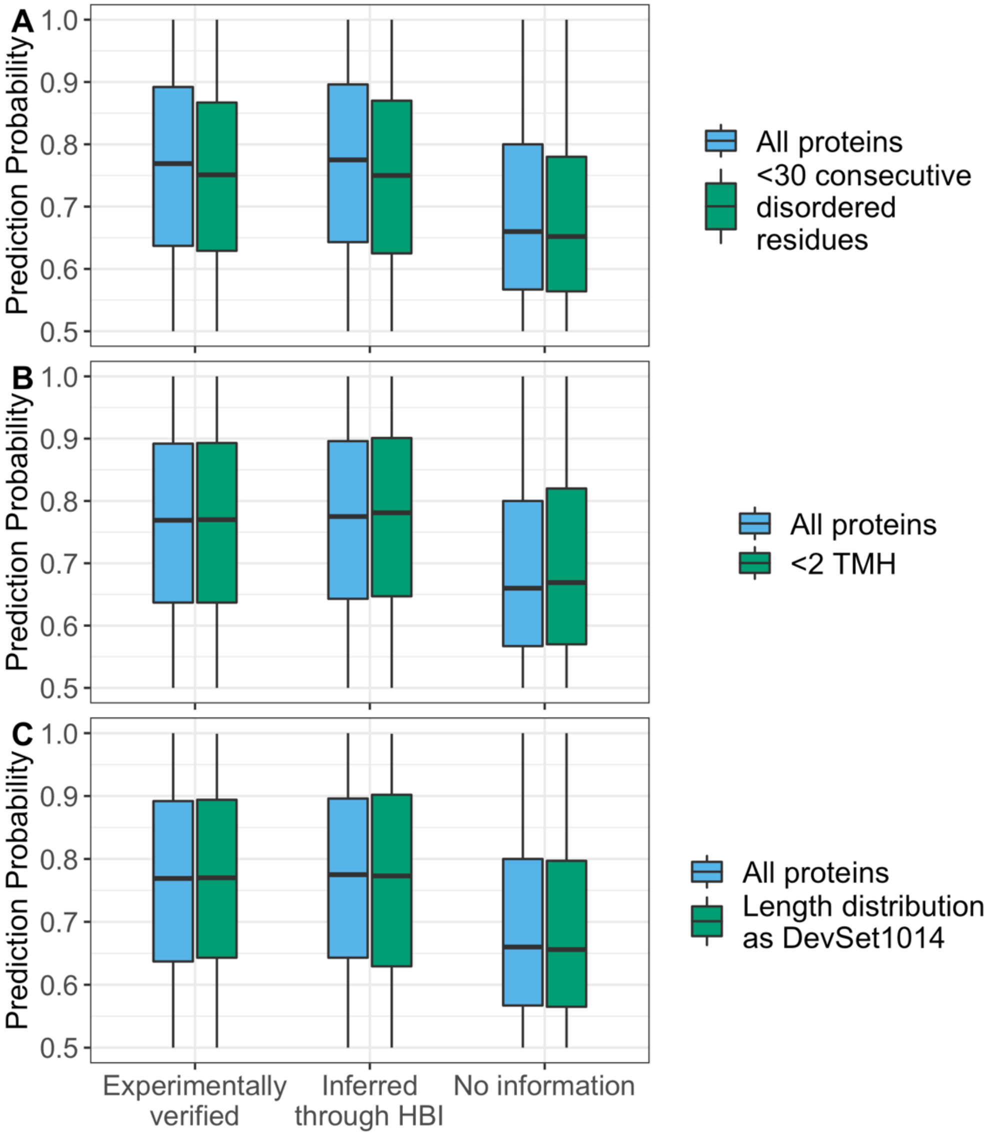

Distribution of prediction probabilities were similar for A. all proteins compared to ordered proteins (<30 consecutive disordered residues; predicted with MetaDisorder[9], B. all proteins compared to non-membrane proteins (< 2 transmembrane helices (TMHs), predicted with TMSEG[10], and C. all proteins compared to proteins of the same length as in the development set DevSet1014. We distinguished 3 subsets of human proteins: Proteins with experimentally verified binding annotations, proteins with binding annotations inferred through homology-based inference (HBI), and proteins without any known binding residues.

### 2. Materials & Methods

#### 2.1. Data sets.

For the construction of our non-redundant data sets, we applied UniqueProt[11] with an HVAL<0. This was a rather strict cutoff which resulted in a reduction of our dataset by more than 90% from 14,894 to 1,314 proteins. To assess whether a less strict cutoff would still lead to a data set of proteins where no pair shares a common binding annotation, we tested how well homology-based inference (HBI) for our training set DevSet1014 (Table S12) would perform using the non-redundant set as lookup set and the HVAL as criterion to decide whether a protein is a homolog or not. Comparing the performance of HBI with our method bindEmbed21DL showed that HBI outperformed bindEmbed21DL for HVAL>0. Only for HVAL=0, performance dropped to the level of bindEmbed21DL. Therefore, the choice of our strict cutoff was necessary to ensure a non-redundant data set although it led to a huge reduction in protein sequences available for training.

---

**Fig. S11: Homology-based inference using HVAL.**

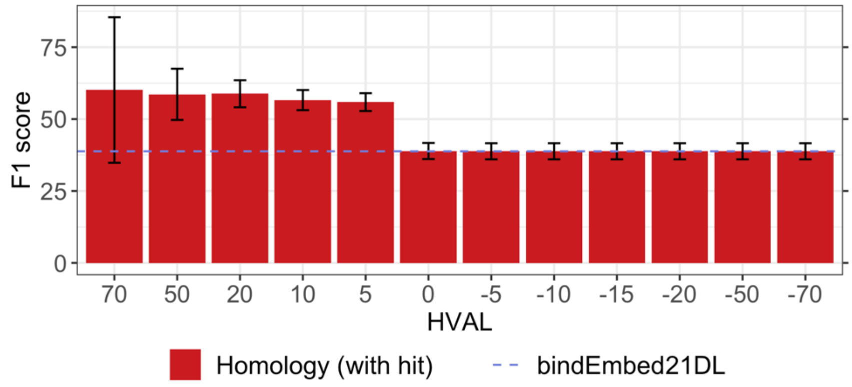

F1 score for homology-based inference using HVAL using DevSet1014 as query set. For HVAL>0, homology-based inference outperformed our Machine Learning method bindEmbed21DL (dashed line). Only when using HVAL=0, performance dropped to the level of bindEmbed21DL. For even lower H-values, no additional hits could be found, probably because no meaningful alignment could be generated. Therefore, performing redundancy reduction at a higher HVAL threshold than zero would lead to a dataset where proteins could share a common binding site.

On the other hand, using even stricter HVAL cutoffs would have led to a tremendous drop in data set size. When reducing our test set TestSet300 at HVAL=-1 against DevSet1014, only 44 proteins remained in the test set. While the number of test proteins dropped further to 11 proteins for an HVAL cutoff of -5, we did not observe any difference in performance (Fig. S12A), indicating that no information leakage appeared for HVAL=0 compared to using even stricter HVAL cutoffs. However, due to the small data set size, confidence intervals were very large for lower HVAL cutoffs. We observed similar results for a redundancy reduction of TestSet300 against DevSet1014 using the E-value: We removed every protein in TestSet300 where we could find a local alignment with a smaller E-value than a certain threshold. For this reduction, the number of proteins was not reduced so largely as for the HVAL cutoff. At E-value=1, our test set still consisted of 202 proteins; for E-value=10, this number dropped to 89 proteins. While the F1 score dropped to 38±5% for E-value=10, it remained within the confidence interval of the performance for the entire set, and performance at lower cutoffs was similar to the overall set (Fig. S12B), again indicating that the redundancy reduction at HVAL=0 yielded a non-redundant data set which did not allow information leakage between train and test set.

---

**Fig. S12: F1 score for TestSet300 redundancy reduced at different HVAL and E-value cutoffs.**

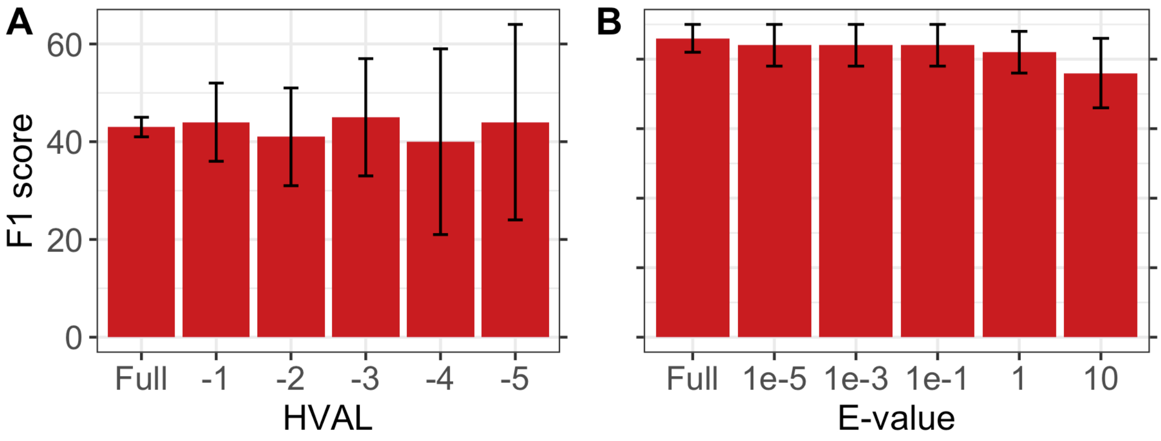

To ensure that our data set split in training (DevSet1014) and test (TestSet300) represented an unbiased split without information leakage between the two independent sets, we assessed performance of further reduced versions of TestSet300 using **A.** stricter HVAL cutoffs and **B.** E-value cutoffs. For both approaches, the F1 score did not change tremendously except for applying an E-value cutoff of 10, where F1 dropped by five percentage points, while remaining within the confidence interval of the performance of the full set.

**Table S12: Development set for bindEmbed21. ***

| | | DevSet1014 | TestSet300 | TestSetNew46 |
|---|---|---|---|---|
| Metal ions | # Proteins | 455 (45%) | 122 (41%) | 15 (33%) |
| | # Binding residues | 2,374 | 881 | 77 |
| | # Non-binding residues | 77,404 | 26,763 | 2,198 |
| Nucleic acids | # Proteins | 108 (11%) | 66 (22%) | 10 (22%) |
| | # Binding residues | 2,689 | 1,470 | 77 |
| | # Non-binding residues | 15,582 | 14,698 | 874 |
| Small molecules | # Proteins | 606 (60%) | 220 (73%) | 25 (54%) |
| | # Binding residues | 9,281 | 3,906 | 425 |
| | # Non-binding residues | 94,119 | 42,629 | 3,269 |
| **All** | **# Proteins** | **1,014** | **300** | **46** |
| | **# Binding residues** | **13,999** | **5,869** | **575** |
| | **# Non-binding residues** | **156,684** | **56,820** | **5,652** |

* The number of proteins, binding residues, and non-binding residues for the three ligand classes (metal ions, nucleic acids, and small molecules) and the three used data sets (DevSet1014, TestSet300, TestSetNew46; see main text for more details). Values from the different ligand classes do not sum to the number for "All" because some proteins are annotated to bind multiple ligands.

#### 2.2. ML architecture

**Fig. S13: Sketch of prediction method.**

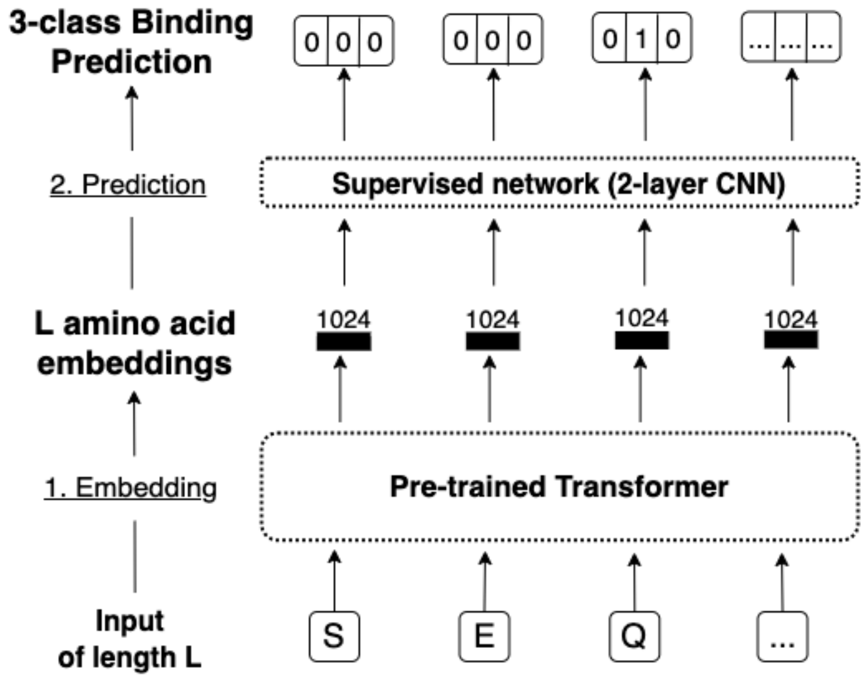

To generate binding residue predictions for three ligand classes, we (1) embed the protein sequence ("SEQ…") using ProtT5[12] to generate 1024-dimensional embeddings for each residue. (2) Those embeddings serve as input for the supervised method consisting of a 2-layer Convolutional Neural Network (CNN). For each residue, the method provides three outputs indicating whether a residue is binding (1) to metal ions, nucleic acids, or small molecules or non-binding (0).

To evaluate whether the fine-grained distinction of three classes of binding residues reduced performance compared to a more coarse-grained binary distinction of binding and non-binding residues, we also trained bindEmbed21DL-binary using the same dataset and architecture as for bindEmbed21DL, but with a dropout rate of 50%, a weight of 4.2 for the positive class (binding residues), and only two output classes to predict whether a residue is binding or non-binding.

#### 2.3. MMseqs2 commands

Our protocol for homology-based inference was entirely based on MMseqs2. To obtain local alignments of query proteins without binding annotations against a set of proteins with known annotations, we performed the following steps and executed the given MMseqs2 commands:

1. Create MMseqs2 database for unlabeled database (used to create profiles), lookup data set with known binding annotations, and query set:
   `mmseqs created in.fasta out.db`

2. Create profiles of query set against large unlabeled database:
   ```
   mmseqs search query.db unlabeled.db result.out tmp/ --num-iterations 2
   mmseqs result2profile query.db unlabeled.db result.out query.profiles
   ```
3. Search profiles against lookup data set:
   ```
   mmseqs search query.profiles lookup.db aln_result_raw.out tmp/ --min-seq-id 0 -s 7.5 – max-seqs 100000 -e 1e-3 -a
   ```
4. Extract local alignments in desired format:
   ```
   mmseqs convertalis query.db lookup.db aln_result_raw.out aln_result.out –format-output "query,target,evalue,nident,mismatch,qstart,tstart,qaln,taln"
   ```

A sample script to perform homology-based inference as implemented for bindEmbed21HBI can also be found on GitHub: https://github.com/Rostlab/bindPredict/blob/master/run_bindEmbed21HBI.py

##### 2.4. Error estimates

We calculated symmetric 95% confidence intervals (CI) assuming a normal distribution of the performance values as error estimates. To investigate whether this could be assumed without affecting the final estimates, we also calculated bootstrapped CIs, i.e., we randomly chose n proteins (n=data set size) with replacement from our data set, calculated the different performance values for each protein and calculated the mean performance over these values. This process was repeated 1,000 times yielding 1,000 mean performances for each metric. Bootstrapped CIs were then calculated as

$$CI_{0.95}(x) = (\bar{x} - t_{0.025} \cdot SD(x), \bar{x} + t_{0.025} \cdot SD(x))$$

where $\bar{x}$ is the average of the 1,000 performance values, $t_{0.025}$ is the Student's t distribution for the confidence level of 95%, and SD is the standard deviation. The resulting performances are given in Table S13. Performance values and error estimates were similar for both calculations of CIs. Therefore, we concluded that a normal distribution can be safely assumed.

**Table S13: Performance estimates for DevSet1014 using CIs and bootstrapped CIs. ***

| Set | | Precision | Recall | F1 | MCC |
|---|---|---|---|---|---|
| DevSet1014 | **CIs** | 37±2% | 52±2% | 39±2% | 0.36±0.02 |
| | **Bootstrapped CIs** | 37±2% | 52±2% | 39±2% | 0.36±0.02 |
| TestSet300 | **CIs** | 46±3% | 52±3% | 43±2% | 0.41±0.02 |
| | **Bootstrapped CIs** | 46±3% | 51±3% | 43±2% | 0.41±0.03 |

* We compared performance calculated using CIs assuming a normal distribution of the per-protein performance values and bootstrapped CIs for the development set (DevSet1014) and the test set (TestSet300). For both sets, we did not observe a difference in the estimated performance.

### 3. Related Work

Many methods focusing on prediction of binding have been reported in the past[13]. However, we did not compare bindEmbed21DL to most of them for various reasons. First, we excluded template-based methods from our comparisons because we see the strength of our method in the area where template-based methods could not be applied (because no template is available). Also, most template-based methods use structural templates and annotations from PDB or BioLiP, i.e., use the annotations from our test set in their template databases. This makes them incomparable to our method because the predictions would just be based on a self-hit of the query protein against its respective template. Secondly, many methods, while focusing on the prediction of binding, do not predict binding residues but rather binding pockets or binding cavities without providing the exact residues involved in binding. Therefore, such methods were not comparable to our approach. Other methods could not be used for comparison because they were simply not available (anymore), or instructions were insufficient for a local installation. Also, other DNA- or RNA-binding prediction methods were excluded because it has been shown that ProNA2020 outperformed its competitors[5]. Table S14 gives a general overview of reasons why methods could not have been used for comparison and lists known binding prediction methods excluded because of those reasons.

**Table S14: Examples of methods not used for comparison and reasons for exclusion. ***

| **Reason for exclusion from comparison** | **Examples of methods** |
|---|---|
| Template-based | GASS-WEB[14], COFACTOR[15], COACH[16], CB-DOCK[17], LIBRA-WA[18], I-LBR[19], COACH-D[20], IonCom[21], 3DLigandSite[22] |
| Prediction of binding pockets or cavities | CSmetaPred[23], CavityPlus[24], DeepSite[25], Kalasanty[26], DeepSurf[27], DeepDrug3D[28], DeepConv-DTI[29], DeepFRI[30], PrankWeb[31] |
| Prediction of catalytic residues | GASS-WEB[14] |
| Not available, installation not possible, or insufficient instructions | FunSite[32], FSCNN[33], CSmetaPred[23], mFASD[34], ZincBinder[35], LigandRFs[36], DeepCSeqSite[37] |

* While many methods focusing on the prediction of binding exist, many could not be compared to our method bindEmbed21DL. Here, we show some example methods and the reasons for not using them for comparison.
